# Lachesin is a mosquito receptor for multiple arthritogenic alphaviruses

**DOI:** 10.64898/2026.07.28.741058

**Authors:** Jesse S. Plung, Enzo Mameli, Wanyu Li, Anja C. M. de Bruin, Jessica A. Plante, Xiaoyi Fan, Biswajit Das, Bailey C. Willett, Yanhui Hu, Ellie M. Hajovsky, Haley Varnum, Praju V. Anekal, Xiaomei Sun, Kate Thornburg, Vesna Brusic, Catherine E. Hammond, Paula Montero Llopis, Raghuvir Viswanatha, W. Robert Shaw, Flaminia Catteruccia, Scott C. Weaver, Kenneth S. Plante, Gisa Gerold, Norbert Perrimon, Jonathan Abraham

## Abstract

Arthritogenic alphaviruses cause acute febrile illnesses associated with rash and arthritis when they are transmitted to humans through the bite of infected mosquitoes. Among these, chikungunya virus (CHIKV), transmitted primarily through the bite of infected *Aedes aegypti* and *Aedes albopictus* mosquitoes, causes explosive outbreaks involving hundreds of thousands to millions of cases, with recent re-emergence in several global regions. While several cellular receptors that mediate alphavirus entry into mammalian cells have been identified, their mosquito counterparts remained unknown, largely due to a lack of functional genomics tools for these invertebrate species. Here, we established a CRISPR-based genetic screening platform in *Aedes albopictus* cells and used it to identify Lachesin, a conserved invertebrate cell adhesion molecule, as a receptor for CHIKV and multiple related alphaviruses including Semliki Forest virus (SFV), o’nyong-nyong virus (ONNV), Mayaro virus (MAYV), and Ross River virus (RRV). Lachesin depletion using RNA interference, anti-Lachesin antibody treatment, and soluble forms of Lachesin blocked CHIKV and SFV E2–E1 glycoprotein-mediated infection of mosquito cells. We show that alphavirus E2–E1 glycoproteins bind the first immunoglobulin domain of Lachesin, facilitating attachment and internalization of virus-like particles. Orthologs from divergent mosquito genera, but not from arachnids or other arthropods, also serve as alphavirus receptors, suggesting that cellular receptor binding is not the main obstacle to arthritogenic alphavirus vector host expansion. Our findings enhance understanding of the mechanisms of alphavirus emergence and vector competence and could aid in the development of broadly active, entry-targeted therapeutics against multiple alphaviruses that threaten public health.

## Introduction

CHIKV is the most widespread arthritogenic alphavirus and causes explosive outbreaks of rash and arthritis in humans. Signs and symptoms of infection include fever, rash, myalgia, and arthralgia, with the latter lasting up to years (*2, 3*). While most human infections tend to be self-limiting, infection can be severe in young children, the elderly, and individuals with medical comorbidities; in certain cases, infection can also be fatal (*4, 5*).

CHIKV is maintained in ancestral, sylvatic, enzootic African cycles involving mammalian reservoir and amplification hosts and forest-dwelling *Aedes* mosquitoes (*6, 7*). However, CHIKV has on multiple occasions emerged into human-amplified, epidemic lineages that have spread throughout Africa, Asia, southern Europe, and, beginning in the 2010s, the Americas (*2, 8-10*), putting 2.8 billion people globally at risk (*11*). In 2024–2025, in addition to re-emergence in endemic regions of the tropics, epidemics have included France, Italy, islands in the Indian Ocean basin, and local transmission was detected in New York State in the United States (*12*). Vector expansion from sylvatic to peridomestic mosquitoes that primarily bite humans, particularly *Ae. aegypti* and *Ae. albopictus*, has propelled CHIKV epidemics (*13-15*).

Additional arthritogenic alphaviruses include ONNV, MAYV, RRV, and Barmah Forest virus (BFV). Other than BFV, they belong to the Semliki Forest serocomplex (SF complex) along with the prototype SF complex alphavirus SFV and the veterinary pathogen Getah virus (GETV), neither of which are associated with arthritis in humans (*13*). Middelburg virus (MIDV), despite its close relationship to the SF complex, has a distinct antigenic profile and is therefore placed in its own serocomplex. BFV belongs to the divergent Barmah Forest serocomplex, but groups with the SF complex within the Old World alphavirus clade. ONNV, MAYV, RRV, and BFV cause locally restricted outbreaks of viral arthritis in various locations around the world, while GETV causes outbreaks among domestic animals widely in Asia, and MIDV causes disease in horses in Central and South Africa. Each virus has been detected in multiple divergent mosquito species, suggesting potential for vector expansion and emergence.

Like most alphaviruses, CHIKV transmission by mosquito vectors is initiated when a viremic bloodmeal contacts the midgut epithelial cells to initiate oral infection. Following replication in the midgut, dissemination to the salivary glands results in replicative shedding into the saliva for transmission during subsequent blood feeding.

To infect mammalian cells, CHIKV can bind the cell adhesion protein matrix remodeling-associated 8 (MXRA8), an immunoglobulin superfamily protein, as a receptor (*16*). MXRA8 has also been reported to serve as a receptor for multiple additional arthritogenic alphaviruses—ONNV, MAYV, RRV, GETV, and BFV (*16*). Additionally, avian and reptilian orthologs of MXRA8 are also receptors for western equine encephalitis virus (WEEV), Sindbis virus (SINV), and a few additional arthritogenic alphaviruses (*17*). However, MXRA8 does not have orthologs in invertebrates including mosquitoes.

Genome-wide screening approaches have only recently become feasible in invertebrate systems (*18*) and, more recently, in disease-vector mosquitoes (*19, 20*), but have not yet been implemented in the major arboviral vector species of the *Aedes* genus (*Culicidae*). Developing these technologies in disease-relevant mosquito species is critical, as vector competence, tissue tropism, and viral adaptation can vary substantially among genera and even among closely related species. Here, building on our previous work (*19*), we establish a CRISPR-Cas9-based functional screening platform in *Ae. albopictus* cells and use it to identify a cellular receptor for CHIKV and additional arthritogenic alphaviruses.

## Results

### Lachesin is a CHIKV and SFV receptor

In previous work, we established a cell line that is derived from *Ae. albopictus* (Asian tiger mosquito) C6/36 cells (*21*) and carries a ΦC31 docking site for library integration (C6/36-HE8) (*19*). Here, to enable pooled CRISPR knockout screening, we added constitutive Cas9 expression to generate a ‘screen-ready’ C6/36-HE8-Ub::Cas9-2A-Neo line (C6/36-HE8C) (Fig. S1A). The alphavirus genome encodes four nonstructural proteins (nsP1–nsP4) and six structural proteins (capsid, E3, E2, 6K, TF, and E1). For the screen, we used a single-cycle reporter virus particle (RVP) system in which the genomic RNA containing the RRV nonstructural proteins, capsid, and a reporter are packaged into virions bearing heterologous alphavirus glycoproteins (*22, 23*). CHIKV contains three major lineages—West African (WA), East/Central/South African (ECSA), and Asian. The Indian Ocean Lineage (IOL) is an ECSA sub-lineage that has caused widespread outbreaks since 2005 (Fig. S2). We performed the screen using RVPs containing the envelope glycoproteins of the WA lineage CHIKV strain 37997 and identified *Lac* (AALFPA_064030, VectorBase) (*24*), which encodes the immunoglobulin superfamily protein Lachesin, as a top hit (Fig. 1A, Table S1).

**Figure 1.**
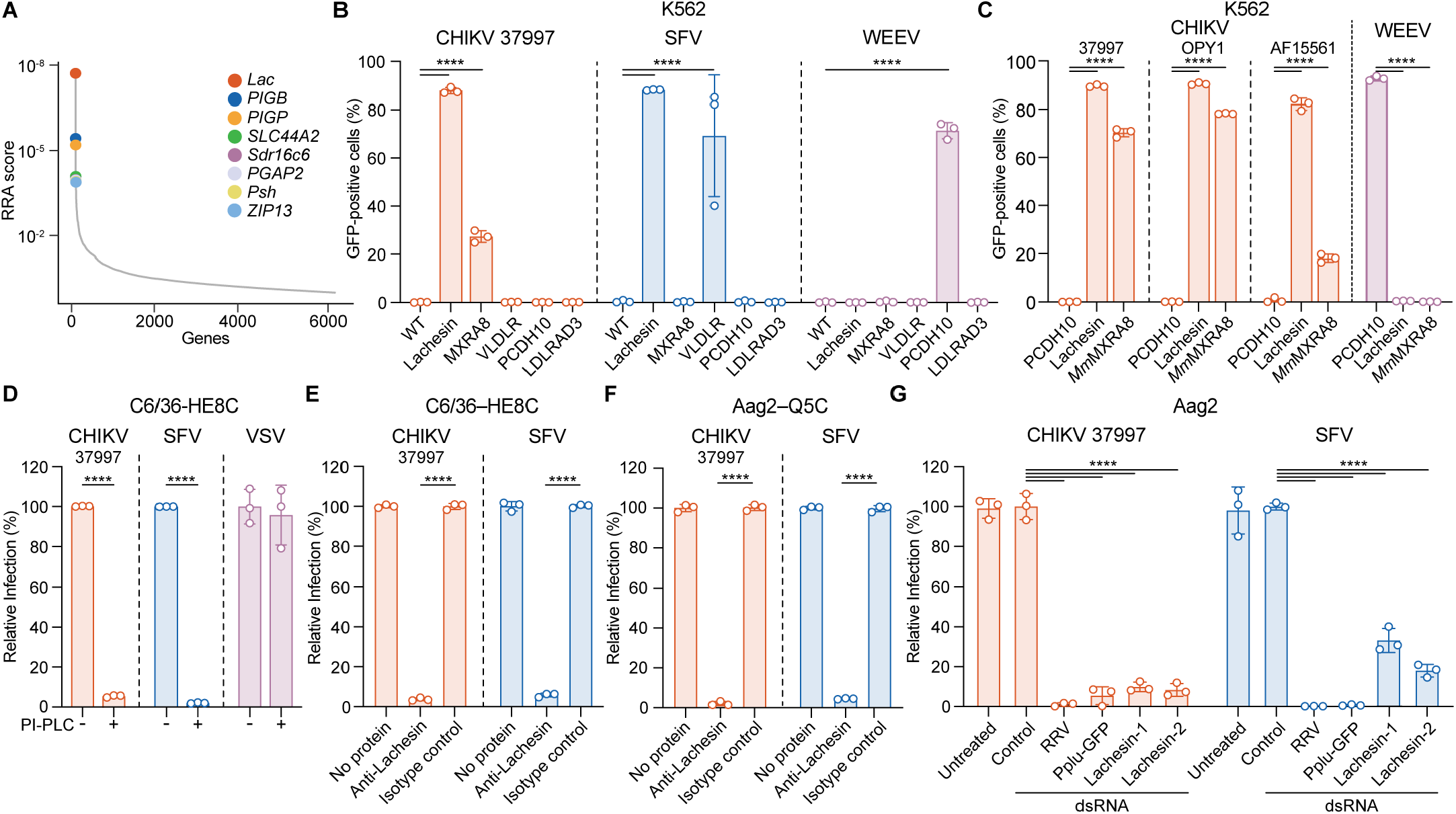
A CRISPR-Cas9 screen identifies Lachesin as a CHIKV and SFV receptor on mosquito cells. **(A)** Results of MAGeCK (*1*) analysis for enriched genes in the CRISPR screen using CHIKV RVP (strain 37997) on *Ae. albopictus* C6/36–HE8C cells, based on top robust rank aggregation (RRA) scores. **(B)** Wild-type (WT) K562 cells or K562 cells stably expressing *Ae. albopictus* Lachesin, or human MXRA8, VLDLR, PCDH10, or LDLRAD3, were infected with GFP-expressing CHIKV (37997), SFV (SFV4), or WEEV (71V1658) RVPs at an MOI of 3 (measured on Vero E6 cells). Infection was monitored by flow cytometry. Cell-surface expression of receptors was confirmed through staining (Fig. 3A–E). **(C)** K562 cells expressing human PCDH10, *Ae. albopictus* Lachesin, or *Mus musculus* MXRA8 were infected with GFP-expressing CHIKV 37997 (WA lineage), OPY1 (ECSA-IOL), or AF15561 (Asian lineage) RVPs, or WEEV (71V1658) RVPs at an MOI of 3 (measured on Vero E6 cells). Infection was monitored by flow cytometry. **(D)** Mock- or phosphoinositide-specific phospholipase C (PI-PLC)-treated *Ae. albopictus* C6/36–HE8C cells were infected with GFP-expressing CHIKV (37997) RVPs, SFV (SFV4) RVPs, or single-cycle vesicular stomatitis virus (VSV) at an MOI of 10 (measured on untreated C6/36– HE8C cells) in the same experiment. Infection was measured by flow cytometry. **(E,F)** C6/36–HE8C cells (**E**) or *Ae. aegypti* Aag2–Q5C (**F**) cells were treated with 10 µg ml^-1^ anti-Lachesin polyclonal antibody or rabbit isotype control polyclonal antibody, followed by infection with CHIKV (37997) or SFV (SFV4) RVPs at an MOI of 1 (measured on C6/36–HE8C cells). Infection was measured by flow cytometry. **(G)** *Ae. aegypti* Aag2 cells were transfected with control double-stranded RNA (dsRNA) targeting LacZ (Control), dsRNA targeting the RVP system including the Ross River virus replicon (RRV) or GFP reporter (Pplu-GFP), or two independent dsRNAs targeting Lachesin. Cells were then infected with CHIKV 37997 or SFV (SFV4) RVPs at an MOI of 1 (measured on C6/36–HE8C cells), with infection measured by GFP expression using flow cytometry. Data are mean ± s.d. of 3 independent experiments performed in duplicate (*n* = 3) (**B**–**F**), or data are mean ± s.d. of 3 independent experiments performed in triplicate (*n* = 3) (**G**). One-way ANOVA with Dunnett’s multiple comparison test, \*\*\*\**P* < 0.0001(**B**–**G**).

Lachesin is a glycosylphosphatidylinositol (GPI)-anchored protein that is involved in cell adhesion and has been implicated in neurogenesis and tracheal system development in insects (*25, 26*). It is highly expressed in the mid-gut of *Ae. aegypti* mosquitoes (Fig. S3A) (*27*), the initial site of infection following a bloodmeal from an infected vertebrate. We also identified three genes that encode proteins involved in GPI anchoring in the screen: *PIGB* (AALFPA_068932), which encodes GPI mannosyltransferase 3, *PIGP* (AALFPA_068174), which encodes phosphatidylinositol N-acetylglucosaminyltransferase subunit P, and *PGAP2* (AALFPA_047349), which encodes post-GPI attachment to proteins factor 2-like (Fig. 1A).

K562 cells are a human lymphoblast-derived cell line that is refractory to entry of all tested alphaviruses; however, they are rendered permissive to infection when transduced to overexpress alphavirus receptors (*22*). We generated a K562 cell line that stably expresses *Ae. albopictus* Lachesin and confirmed cell surface expression using immunostaining (Fig. S4A). Overexpression of *Ae. albopictus* Lachesin or human MXRA8, a known CHIKV receptor (*16*) that served as a positive control, rendered K562 cells permissive to infection by CHIKV 37997 RVPs (Fig. 1B, Fig. S4B). We also included in these assays RVPs for SFV, another alphavirus that belongs to the SF complex and can be vectored by *Ae. aegypti* (*28*). SFV can use human very low-density lipoprotein receptor (VLDLR) and the VLDLR orthologs of *Ae. albopictus* and *Ae. aegypti* to infect cells (*22*). Expression of Lachesin promoted SFV entry into K562 cells, as did overexpression of human VLDLR (Fig. 1B, Fig. S4C). In addition to RVPs for CHIKV 37997 (WA lineage), we also generated GFP-expressing RVPs for representatives of the main epidemic CHIKV lineages, OPY1 (ECSA/IOL lineage) and AF15661 (Asian lineage) and found that these could robustly infect K562 cells expressing *Ae. albopictus* Lachesin and *Mus musculus* MXRA8 (Fig. 1C). *Ae. albopictus* Lachesin overexpression, however, had no effect on infection by RVPs for encephalitic alphaviruses we tested including WEEV, which can use human protocadherin 10 (PCDH10) (*23, 29*), eastern equine encephalitis virus (EEEV), which can use human VLDLR (*22*), and VEEV, which can use human low-density lipoprotein class A domain-containing protein 3 (LDLRAD3) (*30*) (Fig. 1B, Fig. S4F,G). *Ae. albopictus* Lachesin overexpression also did not impact infection by SINV, which can use avian MXRA8 (Fig. S4H) (*17*).

Consistent with the results of experiments with RVPs, K562 cells overexpressing *Ae. albopictus* Lachesin, but not human VLDLR, PCDH10, or LDLRAD3, were rendered permissive to infection by replication-competent CHIKV strains (37997: WA lineage; OPY1: ECSA-IOL lineage) expressing reporters as monitored by flow cytometry (Fig. S5A–E). Additionally, K562 cells expressing Lachesin or human VLDLR were rendered permissive to infection by reporter-expressing, replication-competent SFV (Fig. S5C,F,G) as well as unmodified replication-competent SFV (Fig. S5H).

We next sought to test whether depletion of Lachesin in mosquito cells impairs CHIKV and SFV E2–E1-mediated entry. Lachesin is a GPI anchored protein. Treatment of C6/36-HE8C cells with phospholipase C (PLC), which removes GPI-anchored proteins from the cell surface, decreased cell-surface immunostaining by polyclonal antibody directed against the Lachesin ectodomain and impaired infection of C6/36-HE8C cells by GFP-expressing CHIKV 37997 and SFV RVPs (Fig. 1D, Fig. S6A). The entry of single-cycle vesicular stomatitis virus (VSV) bearing the native VSV glycoprotein (G) and expressing GFP was not affected (Fig. 1D).

Given that *Ae. albopictus* C6/36 cells are deficient in the RNAi pathway (*31*), to determine if the putative mosquito cellular receptor is also used for infection of the other main CHIKV epidemic vector, we also tested *Ae. aegypti* Aag2 cells (*32*) in Lachesin double-stranded RNA (dsRNA)-mediated knockdown experiments. Notably, Lachesin is highly conserved in *Ae. albopictus* and in *Ae. aegypti* (98.5% sequence identity) (Fig. S7A), suggesting that CHIKV would also likely use *Ae. aegypti* Lachesin to infect Aag2 cells. Indeed, knockdown of Lachesin using two independent dsRNA sequences (Lachesin-1 and Lachesin-2), but not a negative control β-galactosidase dsRNA, resulted in decreased infection of Aag2 cells by CHIKV 37997 and SFV RVPs (Fig. 1E, Fig. S6B). Polyclonal antibodies against the ectodomain of Lachesin, but not isotype control antibodies, 37997 and SFV RVP infection in both cell types (Fig. 1F,G, Fig. S6A,C). Collectively, these findings suggest that Lachesin is an important host factor for CHIKV and SFV E2–E1-mediated infection of *Ae. albopictus* and *Ae. aegypti* cells.

### CHIKV and SFV interact with Lachesin

The Lachesin ectodomain contains three immunoglobulin domains (D1, D2 and D3). We generated an Fc fusion protein that contains *Ae. albopictus* Lachesin D1–D3 (Lachesin– Fc) (Fig. S8A). To confirm the interaction of CHIKV and SFV E2–E1 glycoproteins with the Lachesin ectodomain, we transfected HEK 293T cells with plasmids encoding alpha-virus envelope glycoproteins (E3–E2–(6K/TF)–E1) and conducted cell surface staining experiments with Fc fusion proteins. Lachesin–Fc bound cells transfected to express CHIKV 37997 and SFV glycoproteins, but not cells transfected with WEEV (negative control) glycoproteins (Fig. 2A–C). Human (*Hs*) MXRA8-Fc bound cells transfected with CHIKV glycoproteins, but not cells transfected to express SFV or WEEV glycoproteins (Fig. 2A–C, Fig. S8B). We also included as a control an Fc fusion protein that contains the first extracellular cadherin repeat (EC1) of *Passer domesticus* (house sparrow) PCDH10 (*Pd*PCDH10EC1–Fc) (Fig. S8C), which binds with high affinity to WEEV glycoproteins (*33, 34*). *Pd*PCDH10EC1–Fc bound to cells transfected with WEEV glycoproteins but not cells transfected with CHIKV or SFV glycoproteins (Fig. 2A–C). Additionally, in biolayer interferometry experiments, sensor tips coated with Lachesin–Fc, but not a control Fc fusion protein containing human PCDH10 EC1 (*Hs*PCDH10EC1–Fc), bound to CHIKV 37997 and SFV virus-like particles (VLPs) (Fig. 2D,E, Fig. S8D).

**Figure 2.**
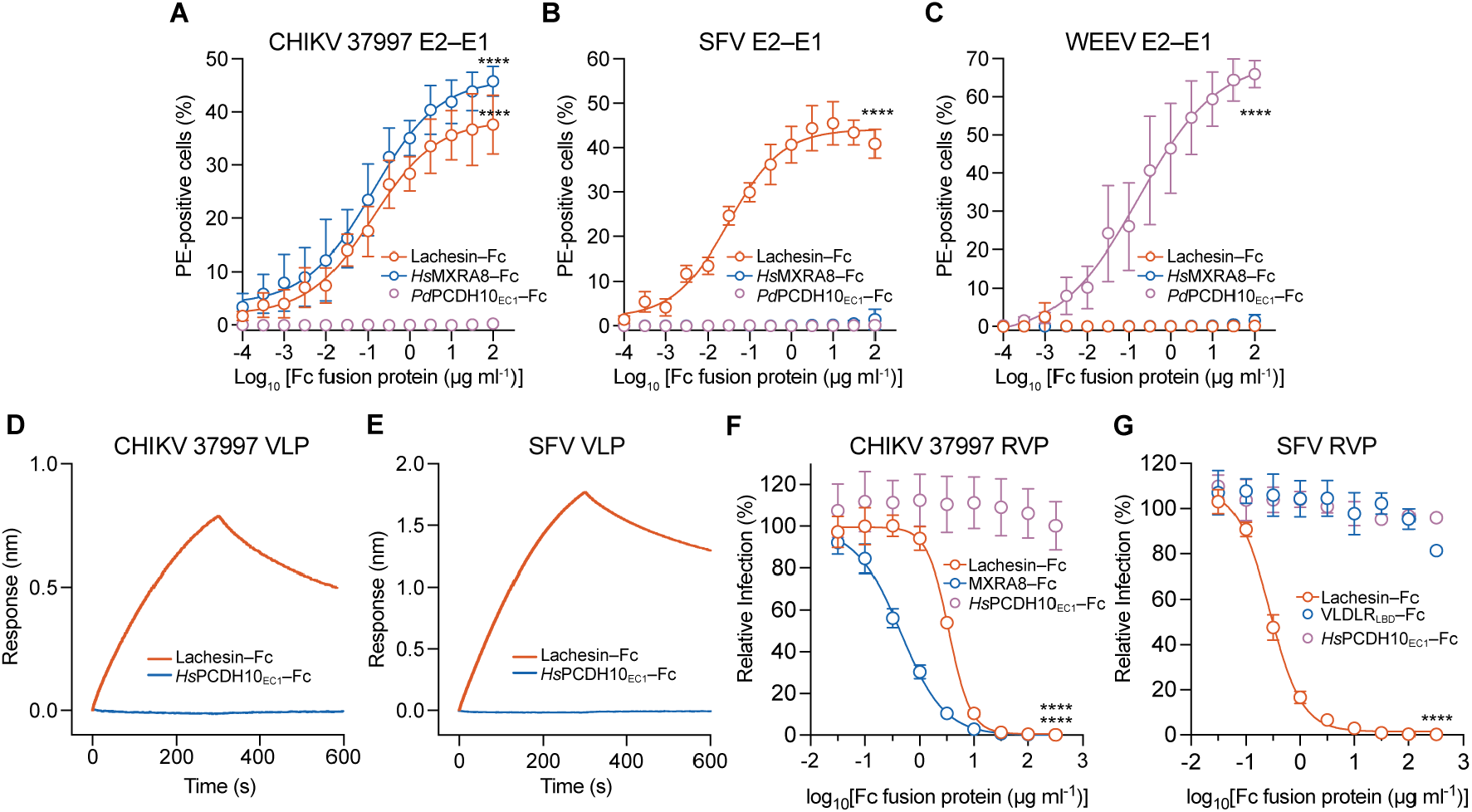
Lachesin interacts with CHIKV and SFV E2–E1 glycoproteins. **(A–C)** Cell surface immunostaining of HEK 293T cells transfected with alphavirus E3–E2–(6K/TF)–E1 proteins of CHIKV (strain 37997) (**A**), SFV (SFV4) (**B**), or WEEV (71V1658) (**C**). Cells were stained with Fc fusion proteins containing the ectodomain of *Ae. albopictus* Lachesin (Lachesin–Fc), the ectodomain of human MXRA8 (*Hs*MXRA8–Fc), or *Passer domesticus* PCDH10 extracellular cadherin repeat 1 (EC1) (*Pd*PCDH10EC1–Fc). PE: *R*-phycoerythrin. **(D–E)** Biolayer interferometry sensorgram of CHIKV (37997) (**d**) or SFV (SFV4) (**e**) virus-like particle (VLP) binding to sensor tips coated with *Ae. albopictus* Lachesin–Fc or an Fc fusion protein containing human PCDH10 EC1 (*Hs*PCDH10EC1–Fc) (control). One representative sensorgram from three experiments is shown for both (**D**) and (**E**). **(F)** *Ae. albopictus* C6/36-HE8C cells were infected with GFP-expressing CHIKV 37997 RVPs at an MOI of 1 (measured on C6/36-HE8C cells) preincubated with the indicated concentrations of Lachesin–Fc, *Hs*MXRA8–Fc, or *Hs*PCDH10EC1–Fc (control). Infection was measured by flow cytometry. **(G)** *Ae. albopictus* C6/36-HE8C cells were infected with GFP-expressing SFV (SFV4) RVPs at an MOI of 1 measured on C6/36-HE8C cells preincubated with the indicated concentrations of Lachesin–Fc, an Fc fusion protein containing the ligand-binding domain (LBD) of human VLDLR (VLDLRLBD–Fc), or *Hs*PCDH10EC1–Fc (control). Infection was measured by flow cytometry. Data are mean ± s.d. from three independent experiments performed in duplicate (*n* = 3) (**A, B, C, F, G**). One-way ANOVA with Dunnett’s multiple comparison test, \*\*\*\**P* < 0.0001 (**A, B, C, F, G**)

### Lachesin–Fc blocks mosquito cell entry

We next tested whether Lachesin–Fc could neutralize CHIKV 37997 and SFV RVP entry into C6/36-HE8C cells. Incubation of RVPs with Lachesin–Fc or *Hs*MXRA8–Fc, but not *Hs*PCDH10EC1–Fc (negative control), blocked CHIKV and SFV RVP entry into *Ae. albopictus* C6/36–HE8C cells (Fig. 2F,G). An Fc fusion protein containing the ligand-binding domain (LBD) of human VLDLR (*Hs*VLDLRLBD–Fc) (Fig. S8E), which binds SFV E2–E1 and can block RVP entry on mammalian cells presumably by blocking access to VLDLR (*22*), did not block SFV RVP entry into C6/36-HE8C cells (Fig. 2G).

### Lachesin D1 is required for entry

We generated a series of Lachesin constructs in which specific domains are deleted, or individual domains are presented (Fig. 3A). We overexpressed these constructs on K562 cells and confirmed their expression using immunostaining against a Flag tag introduced at the N-terminus of each construct (Fig. S6D). A construct in which D3 was deleted (LacΔD3-Flag) robustly supported CHIKV 37997 RVP infection (Fig. 3B). However, a construct in which D1 was deleted (LacΔD1-Flag) did not support CHIKV RVP infection. Overexpression of D1 alone was sufficient to promote CHIKV 37997 and SFV RVP entry, while overexpression of D2 alone did not promote RVP infection. Interestingly, we observed more robust infection of cells overexpressing D1 and D2 (LacΔD3-Flag) than of cells expressing D1 alone. We observed a similar pattern of Lachesin domain dependencies for SFV RVPs, again with D1 being a critical determinant of SFV E2–E1-mediated entry (Fig. 3C). Thus, Lachesin D1 is required and sufficient for CHIKV and SFV E2-E1-mediated entry, but D1 and D2, when combined, more robustly support entry.

**Figure 3.**
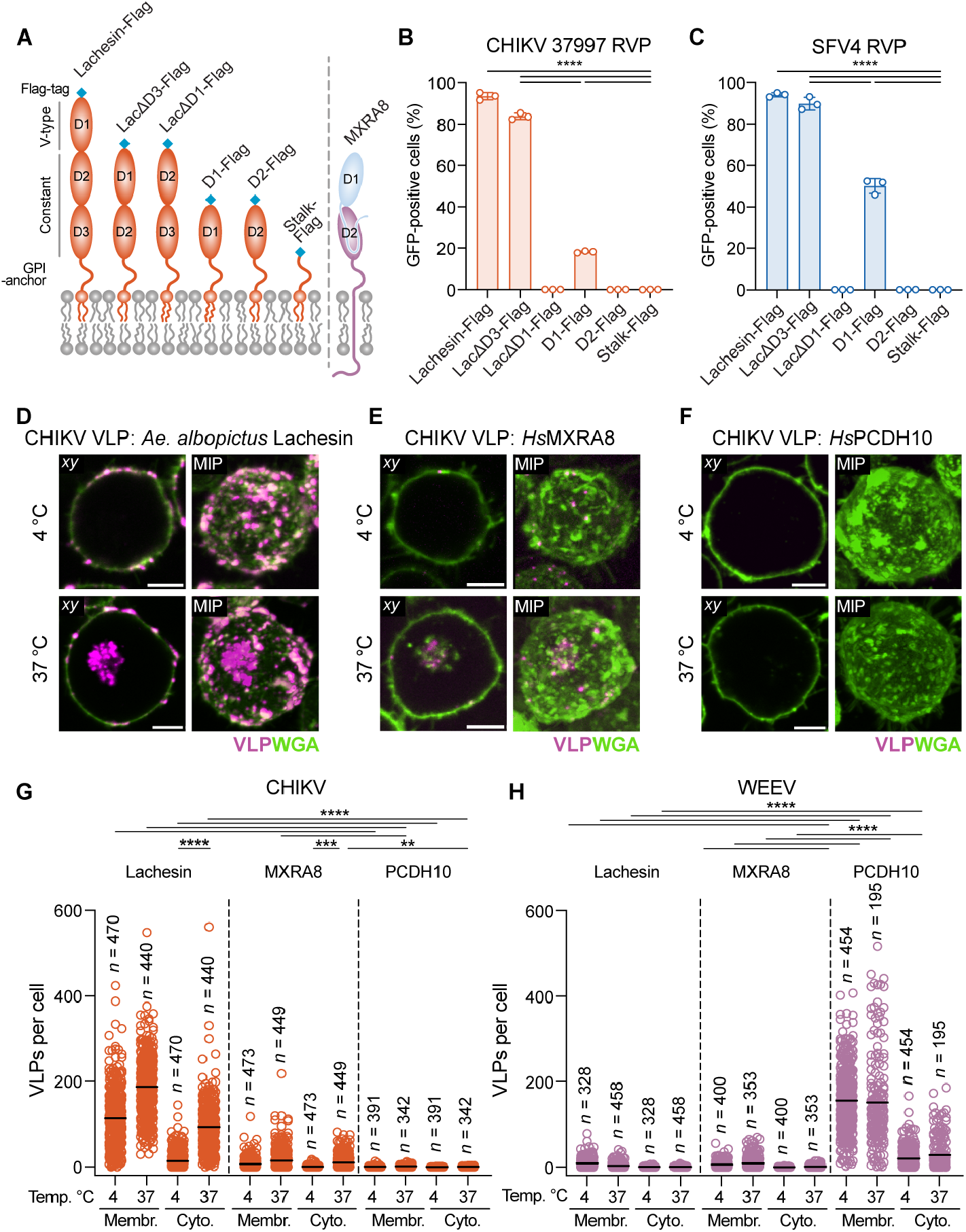
Lachesin immunoglobulin domain 1 mediates attachment and internalization. **(A)** Schematic diagram of Lachesin and MXRA8 domain organization and *Ae*. albopictus Lachesin constructs used for domain mapping analysis. Lachesin immunoglobulin domain types are indicated. **(B,C)** K562 cells stably expressing the Lachesin constructs shown in (**A**) were infected with CHIKV (strain 37997) (**B**) or SFV (SFV4) (**C**), RVPs at an MOI of 3 (measured on K562 cells expressing *Ae. albopictus* Lachesin). Infection was measured using flow cytometry. Cell surface expression of receptors was confirmed through staining (Fig. S6D). **(D–F)**, *xy* slice and maximum intensity projection (MIP) of representative images of wheat germ agglutinin (WGA)-stained K562 cells expressing *Ae. albopictus* Lachesin (**D**), human MXRA8 (**E**), or human PCDH10 (negative control) (**F**) co-incubated with fluorescently labeled CHIKV (strain 37997) VLPs imaged at the indicated temperatures (see Fig. S9, S10 for additional information). Scale bars, 5 μm. **(G,H)** Quantification of CHIKV (37997) (**G**) or WEEV (CBA87) (**H**) VLPs bound to cell membranes (membr.) or internalized into the cytoplasm (cyto.) of individual K562 cells expressing human PCDH10 or MXRA8 at the indicated temperatures (see Fig. S9, S10 for additional information). Data are mean ± s.d. from two experiments performed in triplicates (*n* = 2) (**B, C**). One-way ANOVA Šídák’s multiple comparisons test; \*\*\*\**P* < 0.0001 (**B, C**). Data are means from two experiments with numbers of individual cells indicated; one-way ANOVA with Šídák’s multiple comparisons test; \*\**P* = 0.0012 *\*\*\*P =* 0.0003 \*\*\*\**P* < 0.0001 (**G, H**).

### Lachesin mediates attachment and uptake

We tested whether Lachesin overexpression in K562 cells can support cell surface attachment and internalization of CHIKV VLPs using live-cell confocal microscopy. We fluorescently labeled VLPs for CHIKV 37997 and WEEV (Fig. S9A) and incubated these with K562 cells overexpressing *Ae. albopictus* Lachesin, human MXRA8, or human PCDH10. Lachesin expression allowed CHIKV binding to cell surface membranes (Fig. 3D–G, Fig. S9B, S10). An increase in particles was detected in the cytoplasm of cells expressing Lachesin at 4 °C versus 37 °C, suggesting internalization. An increase in CHIKV VLPs in the cytoplasm of cells expressing MXRA8 (positive control) but not PCDH10 (negative control) was also detected at 4 °C versus 37 °C (Fig. 3D–G). Conversely, ectopic expression of PCHD10, but not Lachesin or MXRA8, allowed for WEEV VLP binding and internalization (Fig. 3H, Fig. S9B). Thus, Lachesin can bind CHIKV E2–E1 to facilitate particle cell surface binding and internalization.

### Lachesin recognition by alphaviruses

Other than CHIKV and SFV the SF complex includes the human arthritogenic viruses ONNV, RRV, MAYV, the veterinary pathogen GETV, and two viruses currently not known to cause diseases in humans and animals, Bebaru (BEBV) and Una (UNAV) (Fig. 4A). The SF complex, together with Middelburg (MIDV), Ndumu (NDUV), and Barmah Forest (BFV) viruses are within the Old World alphavirus clade based on E2–E1 glycoprotein phylogenetic analysis. (Fig. 4A). Despite their genetic diversity, alphaviruses are routinely isolated and propagated on C6/36 cells (*35-45*). The broad permissiveness of these cells to productive infection suggests the presence of receptors on this *Ae. albopictus* cell line for many alphaviruses.

**Figure 4.**
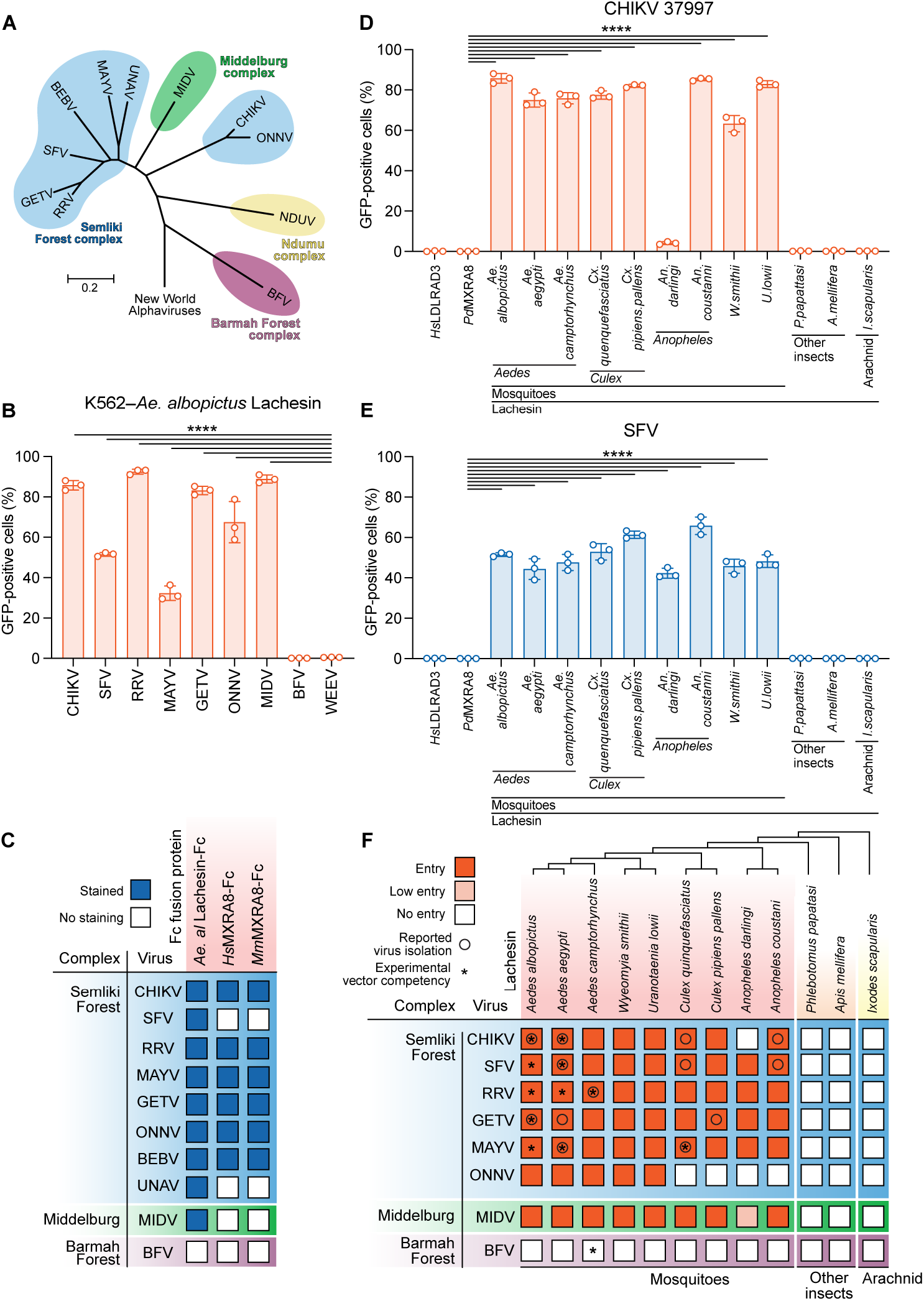
Multiple alphaviruses use diverse lachesin orthologs. **(A)** Phylogenetic tree of Semliki Forest, Middelburg, Ndumu, and Barmah Forest complex alphaviruses based on sequence alignment of their E2 glycoproteins. New World alphaviruses: WEEV, VEEV, EEEV, and SINV were used as an outgroup (not shown). **(B)** K562 cells stably expressing *Ae. albopictus* Lachesin were infected with the indicated GFP-expressing RVPs with infection measured using flow cytometry. Strains: CHIKV 37997; SFV SFV4; RRV T48; MAYV LET-1430; GETV M1; ONNV SG650; MIDV MIDV857; BFV K61404; WEEV McMillan. **(C)** Summary of the results of cell surface staining of K562 cells expressing E2–E1 glycoproteins of the indicated alphaviruses, stained with *Ae. albopictus* Lachesin–Fc, human MXRA8–Fc, or murine (*Mm*) MXRA8ect–Fc. See Fig. S12 for additional information. **(D,E)** K562 cells expressing Lachesin orthologs were infected with GFP-expressing CHIKV 37997 (**D**) or SFV4 (**E**) RVPs at an MOI of 2 (measured on Vero E6 cells). Infection was measured using flow cytometry 24 h post-infection. **(F)** Summary of the results of alphavirus RVP infectivity assays with K562 cells overexpressing the indicated Lachesin orthologs. A toggled phylogenetic tree of Lachesin orthologs is shown (See Fig. S7B for more information). “Entry” signifies more than 10% infection as assessed GFP positive cells, and “Low entry” signifies 5–10% GFP positive cells in the assay. Asterisks indicate that a given virus has been isolated from that mosquito species. See Fig. S12 for additional information. Data are mean ± s.d. of 3 independent experiments performed in duplicate (*n* = 3); one-way ANOVA with Dunnett’s multiple comparisons test *\*\*\*\*P* < 0.0001 (**B, D, E**)

We tested whether additional SF, MID, and BF complex alphaviruses could use Lachesin as a receptor. K562 cells overexpressing *Ae. albopictus* Lachesin were rendered permissive to infection by RVPs bearing the E2–E1 glycoproteins of the SF complex members CHIKV, SFV, RRV, MAYV, GETV, and ONNV, and the non-SF complex Old World alphavirus MIDV, but not BFV (Fig. 4B). Additionally, replication-competent RRV could infect K562 cells overexpressing *Ae. albopictus* Lachesin (Fig. S5C,I,J). Consistent with these infectivity assays, *Ae. albopictus* Lachesin– Fc could stain HEK 293T cells transfected to express the glycoproteins of RRV, MAYV, GETV, ONNV, and MIDV, but not BFV (Fig. 4C, Fig. S11A).

We were unable to generate RVPs for SF complex viruses BEBV and UNAV to evaluate their ability to engage overexpressed Lachesin to infect K562 cells. However, Lachesin– Fc stained HEK 293T cells transfected to express the BEBV and UNAV glycoproteins, (Fig 4C, Fig. S11A), suggesting that the E2–E1 glycoproteins of these viruses can also bind *Ae. albopictus* Lachesin. Collectively, these findings suggest that *Ae. albopictus* Lachesin can broadly serve as a cellular receptor for SF and MID complex alphaviruses.

### CHIKV recognition of Lachesin orthologs

While CHIKV is principally vectored by *Ae. aegypti* and *Ae. albopictus* during epidemics, CHIKV has been reportedly isolated from diverse mosquitoes of the *Culex, Coquillettidia, Wyeomyia*, and *Mansonia* genera (*15, 46*), which together with the *Aedes* genus, fall under the subfamily *Culicinae*. CHIKV has also been isolated from *Anopheles* mosquitoes (*15*), which belong to the divergent mosquito subfamily *Anophelinae*, estimated to have diverged from *Culicinae* approximately 179 million years ago (*47*).

To further examine the potentially broad usage of this putative receptor, we selected a wide range of insect Lachesin protein sequences to study their ability to facilitate CHIKV E2–E1-mediated entry (Fig. S7A). For *Aedes* spp., we used Lachesin sequences for two CHIKV vectors, *Ae. albopictus* and *Ae. aegypti*, and *Ae. camptorhynchus* (southern saltmarsh mosquito), which is a vector for RRV and BFV in Australia. We also included two mosquito species that mostly do not bite mammals, *Wyeomyia smithii* (pitcher plant mosquito) and *Uranotaenia lowii* (pale-footed Uranotaenia), because they are grouped closely with *Aedes* lachesin orthologs (Fig. S7B). For *Culex* spp., we included the orthologs of *Cx. quinquefasciatus* (southern house mosquito) and *Cx. pipiens pallens* (northern house mosquito). For *Anopheles* spp., we included *An. darlingi* and *An. coustani*, which are known malaria vectors. Lastly, we also included orthologs from two non-mosquito insects, *Phlebotomus papatasi* (sandfly), *Apis mellifera* (honeybee), and the ortholog of the arachnid *Ixodes scapularis* (deer tick). We confirmed expression of these proteins on the surface K562 cells using immunostaining (Fig. S7C). Sequence analysis demonstrates a high level of sequence conservation of this protein among widely divergent mosquitoes, and phylogenetic analysis is largely congruent with the phylogeny of mosquito genera and subfamilies (*47*).

CHIKV 37997 RVPs could infect K562 cells overexpressing Lachesin orthologs of most tested mosquito species (Fig. 4D), except for the *An. darlingi* ortholog, for which we only observed low levels of infection. CHIKV RVPs recognized Lachesin orthologs of multiple mosquitoes from which CHIKV isolation has been reported, including its other principal epizootic vector, *Ae. aegypti*, consistent with the results of Lachesin dsRNA knockdown and antibody blocking studies performed with *Ae. aegypti* Aag2 cells (Fig. 1E,G, Fig. 4D). CHIKV RVPs could not infect K562 cells expressing Lachesin orthologs of non-mosquito arthropods and the tick ortholog we tested.

### Broad viral recognition of mosquito Lachesin

Like CHIKV, many arthritogenic alphaviruses have been isolated from a wide array of mosquitoes. Accordingly, we found that RVPs for SFV, RRV, MAYV, GETV, ONNV, and MIDV could infect K562 cells expressing Lachesin orthologs from divergent mosquito genera, including reported sources of natural isolation for these viruses (Fig. 4E, F, Fig. S12). Surprisingly, although ONNV is the only pathogenic alphavirus known to use *Anopheles* spp. mosquitoes as principal vectors, ONNV RVPs could not recognize tested *Anopheles* Lachesin orthologs that were otherwise recognized by RVPs for CHIKV, SFV, RRV, MAYV, GETV, and MIDV (Fig. 4F, Fig. S12). As with CHIKV RVPs, those for the additional alphaviruses we tested could not infect K562 cells overexpressing non-mosquito Lachesin orthologs (Fig. 4D–F, Fig. S12).

BFV could not infect K562 cells expressing any of the Lachesin orthologs we tested, including that of its natural vector, *Ae. camptorhynchus* (southern saltmarsh mosquito). Interestingly, while both RRV and BFV are naturally vectored by *Ae. camptorhynchus*, RRV but not BFV RVPs could recognize *Ae. camptorhynchus* Lachesin to infect K562 cells (Fig 4E, Fig. S12), suggesting that this mosquito expresses different, BFV-specific receptors.

## Discussion

The emergence of large-scale outbreaks of CHIKV is due in part to increased adaption to the peridomestic vector, *Ae. albopictus*. A single amino acid substitution on CHIKV E1— A226V—a hallmark of the IOL sub-lineage, has been shown to increase *Ae. albopictus* vector competence (*48, 49*). The CHIKV strains we tested containing the native E1 residue A226 (strain 37997 from the WA lineage, and AF15561 from the Asian lineage) or the V226 polymorphism (OPY1, an ECSA-IOL strain) (Fig. S2) were able to engage *Ae. albopictus* Lachesin (Fig. 1C). The effect of this polymorphism on viral fitness in mosquitoes may thus act downstream of receptor engagement, which is consistent with prior studies suggesting that the A226V substitution modulates pH and cholesterol dependence during viral entry (*48, 50*).

CHIKV has been detected in mosquito genera that are not recognized as primary vectors; however, these detections may represent residual viral presence in a recently ingested blood meal and not productive infection, or an infection limited to the midgut and not disseminated to the hemocoel for infection of the salivary glands. Indeed, CHIKV has been detected in *Cx. quinquefasciatus, An. gambiae*, and *An. funestus*, but these mosquitoes appear refractory to productive CHIKV infection (*15, 51-54*). These vector incompatibilities may arise from barriers at the midgut or other tissues due to a lack of tissue-or species-specific host factors, whose absence could potentially obstruct various stages of the CHIKV replication cycle (*55, 56*). Our results will facilitate a molecular dissection of barriers to vector competence. For example, we found that CHIKV can recognize *Cx. quinquefasciatus* Lachesin, suggesting that receptor binding is not a barrier to infection in this CHIKV-refractory mosquito. This may be the case for other mosquito species that are resistant to CHIKV but express Lachesin orthologs to which CHIKV can bind. Future studies examining the tissue distribution of Lachesin in divergent mosquito species may aid in the interpretation of mosquito susceptibility to CHIKV infection.

We also observed evidence of species-specific recognition of *Anopheles* Lachesin orthologs by CHIKV RVPs. CHIKV is closely related to ONNV, but whereas ONNV is transmitted by *Anopheles* mosquitoes in nature, CHIKV is not (*53*). However, chimeric ONNV bearing CHIKV structural proteins efficiently infects the ONNV vector *An. gambiae* to comparable levels as wild-type ONNV, and the molecular determinants of ONNV vector specificity are thought to reside in nsP3 (*57*), suggesting that the barrier to CHIKV replication in certain *Anopheles* mosquitoes occurs downstream of viral entry. Furthermore, *An. stephensi*, while not thought to be a vector of CHIKV in nature, is able to transmit CHIKV in laboratory settings (*58, 59*). ONNV RVPs efficiently recognized tested *Aedes* Lachesin orthologs, but not *Anopheles* Lachesin orthologs. It is possible that ONNV recognizes Lachesin orthologs of its natural vectors *An. gambiae* and *An. funestus* but not of the two species we tested. Alternatively, ONNV may have evolved to recognize distinct receptors in *Anopheles* vectors.

Multiple SF complex alphaviruses including SFV, RRV, MAYV, and GETV have been isolated from atypical mosquito species other than their primary vectors (*60-64*). Laboratory vector competence experiments are needed to discern incidental viral presence in blood meals from productive infection as the cause of virus detection during surveillance. Such studies revealed mosquitoes from divergent genera can serve as competent vectors for these viruses in a laboratory setting (*38, 65-68*). The ability of the E2–E1 proteins of these alphaviruses to broadly recognize Lachesin orthologs could at least in part underlie these observations.

Lachesin is highly expressed in the midgut of adult *Ae. aegypti* mosquitoes based on the results of prior single-cell RNA sequencing experiments (Fig. S3) (*27, 69*), and would thus be positioned at site of initial oral mosquito infection following an infectious blood meal. In addition to expression in the gut, Lachesin is also expressed in the salivary glands, suggesting that it may play a role in transmission. Additional studies will be required to explore the tissue-specific role of Lachesin as a cellular receptor in mosquitoes.

Although Lachesin lacks a transmembrane anchor and cytoplasmic tail to mediate virus internalization, prior work suggests that endocytosis of GPI-anchored proteins can nonetheless occur through clathrin-dependent and - independent pathways (*70-72*). Interestingly, although extensively studied mammalian alphavirus receptors— MXRA8, LDLRAD3, VLDLR, and PCDH10—contain TM anchors, replacing these TM anchors with GPI anchors still allows these molecules to serve as functional receptors (*16, 23, 30*).

Lachesin does not serve as a receptor for several encephalitic alphaviruses (EEEV, WEEV, and VEEV) or SINV and is not recognized by the divergent arthritogenic alphavirus BFV. These observations suggest that Lachesin recognition is an evolutionary innovation of the SF and MID complexes, and that BFV and New World, encephalitic alphaviruses likely evolved to bind distinct mosquito vector receptors.

Given that CHIKV must infect mammalian amplification hosts and mosquito vectors sequentially to complete its transmission cycle, its glycoproteins may be under balancing selection to maintain affinity for both MXRA8 and Lachesin. Additional studies will be required to examine the mutational constraints on alphavirus E2–E1 glycoprotein residues that contact both mammalian and mosquito receptors.

While MXRA8 was implicated in BFV infection based on replication assays conducted in Mxra8 deficient murine fibroblasts (3T3 cells) (*16*), we did not detect interaction between cells expressing BFV glycoproteins and murine or human MXRA8–Fc (Fig. 4C, Fig. S8B,F, S11B–D). Differences in the BFV strain sequences (we used BFV strain K61404 and Zhang et al. used strain K10521) could potentially account for this discrepancy. UNAV replication in murine 3T3 cells was not affected by Mxra8 genetic knockout (*16*), consistent with the lack of murine and human MXRA8–Fc staining of cells expressing UNAV glycoproteins (Fig. 4C, Fig. S11B,C).

Overall, our work suggests that Lachesin is an alphavirus receptor that promotes mosquito cell infection by multiple human and veterinary pathogens in the SF and MID complexes. Based on our observation of promiscuous viral recognition of mosquito Lachesin orthologs, we hypothesize that a weak barrier at the level of cellular receptor binding could allow arthritogenic alphaviruses to explore and adapt to new mosquito vector species for transmission.

## Acknowledgements

This work was supported by Burroughs Wellcome awards to J.A., a Vallee Scholar award to J.A., a Smith Family Foundation Odyssey award to J.A., a Charles E.W. Grinnell Trust award to J.A., NIH award R01 AI182377 to J.A., and a G. Harold and Leila Y. Mathers Foundation award to J.A. This work was also supported by the UTMB Institute for Human Infections and Immunity award to K.S.P. and by NIH awards T32AI700245 to J.S.P., T32GM008313 and T32GM144273 to H.V., and R24AI120942 to S.C.W. Work in the Perrimon laboratory is supported by NIH 5P41GM132087, and 3R01AI170835. This work was also supported by a Research Grant from HFSP (RGP011/2023 award DOI 10.52044) to G.G. and N.P.. We thank Tonya M. Colpitts (Boston University, NEIDL) for providing the *Ae. aegypti* Aag2 cell line. The authors acknowledge the MicRoN (Microscopy Resources on the North Quad) Core for help with live-cell imaging, Biosynth International Ltd. for help with antibody generation, and the DRSC-FGR (Drosophila RNAi Screening Center-Functional Genomic Resources) facility at Harvard Medical School for RNAi reagents. F.C., N.P., and J.A. are investigators of the Howard Hughes Medical Institute. We thank Bridget Golan for assistance with illustration editing.

## Author contributions

E.M. designed the sgRNA library and screening cell line with the help of R.V. and Y.H. J.S.P. performed the CRISPR-Cas9 genetic screen with assistance from E.M. E.M. performed the dsRNA knockdown experiments and infectivity studies, with assistance from Y.H., X.S., and R.V. J.S.P. and W.L. generated human cell lines expressing receptors, RVPs, and recombinant proteins, and performed infectivity studies and confocal microscopy experiments, with the help of X.F., H.V., C.E.H., and V.B.. W.L. and J.S.P. performed phylogenetic analysis. A.C.M.d.B. conducted infectivity studies with replication-competent alphaviruses expressing reporter proteins rescued from molecular clones, and G.G. supervised those studies. X.F. produced VLPs and conducted biolayer interferometry experiments. J.A.P. designed and conducted infectivity studies with replication-competent SFV rescued from a molecular clone with the help of E.M.H., and S.C.W. and K.S.P. supervised those experiments. Y.H. contributed to sgRNA library design. P.V.A. and P.M.L. helped conduct and analyze live-cell confocal microscopy studies. B.D., B.C.W., K.T., W.R.S., and F.C. conducted experiments. N.P. and J.A. supervised the study, and S.C.W., K.S.P., N.P., and J.A. acquired funding. J.S.P. and W.L. wrote the original draft of the manuscript, and all authors participated in reviewing and editing.

## Materials and Methods

### Cells and viruses

HEK293T cells (human kidney epithelial, ATCC CRL-11268) and Vero E6 cells (*Cercopithecus aethiops* kidney, ATCC CCL-81) were maintained in Dulbecco’s modified Eagle’s medium (DMEM, Gibco) supplemented with 10% (v/v) fetal bovine serum (FBS) and 25 mM HEPES (Thermo Fisher Scientific) for titration of reporter viruses or with 10% (v/v) FBS and 1% (v/v) penicillin-streptomycin for work with SFV A774wt. K562 (human chronic myelogenous leukemia, ATCC CCL-243) cells were maintained in RPMI1640 (Thermo Fisher Scientific) supplemented with 10% (v/v) FBS, 25 mM HEPES, and 1% (v/v) penicillin-streptomycin. Expi293F™ cells (Thermo Fisher Scientific A14527) were maintained in Expi293 Expression Medium (Thermo Fisher Scientific). Cell lines were not authenticated. Absence of mycoplasma is confirmed through routine mycoplasma test using e-Myco PCR detection kit (Bulldog Bio 25234). BHK-21 [C-13] cells (Syrian golden hamster fibroblast, ATCC CCL-10) and BHK-21/WI2 (Syrian golden hamster fibroblast, Kerafast EH1011) were cultured in Eagle’s Minimum Essential medium (EMEM, Gibco) supplemented with 10% (v/v) FBS and 25 mM HEPES (Thermo Fisher Scientific).

The *Ae. albopictus* clonal cell line C6/36–HE8 (*19*) (DGRC Stock 336; RRID:CVCL_B3N5), originally derived in the Perrimon laboratory, was maintained in Schneider’s insect medium (Gibco) supplemented with 10% (v/v) heat-inactivated FBS (Gibco) and 1% (v/v) penicillin-streptomycin (Gibco). In some experiments this standard media formulation was supplemented with 25 mM HEPES, 1× MEM non-essential amino acids (Gibco). To generate a Cas9-expressing derivative, C6/36-HE8 cells were stably transfected with the plasmid pAaePUb::Cas9-2A-Neo (GenBank OL312705; Addgene plasmid #176680)(*19*) and selected for 30 d in medium containing 500 µg ml^-1^ Geneticin (G418 sulfate; GoldBio). The resulting line, designated C6/36-HE8-Ub::Cas9-2A-Neo (attP_+_ Cas9_+_; RRID pending registration and deposit), was maintained in Geneticin-supplemented medium and used as the “screen-ready” C6/36–HE8C *Ae. albopictus* line for both the CRISPR knockout screening and infection-related experiments in this study. The *Ae. aegypti* cell line Aag2 (RRID:CVCL_Z617)(*32*), kindly provided by Tonya M. Colpitts (Boston University, NEIDL) was maintained in Schneider’s insect medium supplemented with 10% (v/v) heat-inactivated FBS and 1% (v/v) penicillin-streptomycin, or alternatively for RNAi experiments in Leibovitz’s L-15 medium containing 10% (v/v) FBS, 1% (v/v) tryptose phosphate broth (Gibco), and 1% (v/v) penicillin-streptomycin. For in vitro antibody blocking experiments, we used the clonal cell line Aag2-Q5-Ub::Cas9-2A-Neo (referred as Aag2–Q5C throughout the manuscript) derived from native Aag2 cells following a previously established protocol (*19*). Briefly, Aag2 native cell line were co-transfected using Effectene (Qiagen) and a mixture of plasmids encoding HSP70-Minos transposase and three separate MiMIC mCherry plasmids for frame selection with molar ratio of 1:0.3:0.3:0.3 respectively. After transfection, cells were passaged three times, expanded, and single-cell sorted by fluorescence-activated cell sorting (FACS) into 96-well plates to obtain clonal mCherry-trapped populations. FACS was performed using a MoFlo Astrios (BD) with a 100 µm nozzle at 20 psi (Harvard Medical School, Immunology Flow Core). A selected Minos-transformed, mCherry positive, clonal cell line was established, herein named Aag2-Q5. Aag2-Q5 clone was subsequently transfected with pAaePUb::Cas9-2A-Neo (GenBank OL312705; Addgene plasmid #176680) (*19*) and selected in 200 µg ml^-1^ Geneticin (Gibco) for 30 d to obtain Aag2Q5-Ub::Cas9-2A-Neo. We note that the Aag2-Q5 clone and derived Aag2– Q5C cell line have lost mCherry signal as a result of potential silencing, and RMCE cassette presence and functionality is unverified. All mosquito cell culture, transfection, and infections were performed at 25 °C or 27 °C

Virus stocks of infectious clones from CHIKV OPY1 5′ GFP and CHIKV 37997 5′GFP (kind gifts from G. Simmons71,72), and VEEV TC-83 GFP (kind gift from I. Frolov73) expressing GFP under the control of a subgenomic promoter, and SFV4-GFP (kind gift from A. Merits74) and RRV T48-mCherry (kind gift from R. Kuhn, C. Schmidt, and B. Schnierle75) harboring GFP and mCherry fused to nsP3 respectively, were generated upon electroporation of in vitro transcribed mRNA or plasmid DNA in BHK-21 cells76. Tissue Culture Infectious Dose 50% (TCID50) titers were determined by endpoint titration in BHK-21 cells. Titers for SFV A774wt were determined by plaque assay on Vero cells.

### Reporter virus particle generation

To generate reporter virus particles (RVPs), HEK 293T cells were transfected with two plasmids using Lipofectamine 3000 (Thermo Fisher Scientific) by following the manufacturer’s instructions. One plasmid is a modified pRR64 RRV replicon (*73*) provided by R. Kuhn (Purdue University) in which the Sp6 promoter was exchanged with a CMV promoter, and the E3–E2–(6K/TF)–E1 sequence was replaced with a GFP or GFP-CD20 reporter preceded by a porcine teschovirus-1 2A self-cleaving peptide. The second plasmid is a pCAGGS or pTWIST-CMV-BG-WPRE-Neo expression vector containing the E3–E2–(6K/TF)–E1 sequence of heterologous alphaviruses. At 4–6 h post transfection, media was replaced with Opti-MEM supplemented with 5% (v/v) FBS, 25 mM HEPES (Thermo Fisher Scientific) and 5 mM Sodium Butyrate. Two days after transfection, supernatant was harvested and spun down at 3000g for 5 min before being passed through a 0.45 µm filter, aliquoted and stored at -80 °C.

Alphavirus E3–E2–(6K/TF)–E1 sequences included in the pCAGGS vectors were as follows: CHIKV strain 37997 (GenBank AY726732), CHIKV strain OPY1 (GenBank KT449801), SFV strain SFV4 (GenBank AKC01668), MAYV strain LET-1430 (GenBank PP505832), WEEV strain 71V1658 (GenBank GQ287645), WEEV strain McMillan (GenBank GQ287640), EEEV strain FL91-469 (GenBank AY705241), and VEEV strain INH-9813 (GenBank KP282671). Alphavirus E3–E2–(6K/TF)– E1 sequences included in the pTwist-CMV-BG-WPRE-Neo vectors were as follows: CHIKV strain AF15661 (GenBank MK028840), ONNV strain SG650 (GenBank YP010775618), RRV strain T48 (GenBank ACV67002), GETV strain M1 (GenBank EU015061), BFV strain K61404 (GenBank MN689027), and MIDV strain MIDV857 (GenBank EF536323). For a list of all virus sequences used in this study, see Table S2.

### Reporter virus particle titration

GFP-expressing RVPs were titered on Vero E6 cells, C6/36–HE8C cells, and K562 cells overexpressing *Ae. albopictus* Lachesin seeded in 96 well plates using tenfold serial dilutions of RVP stocks. At 24 h post-infection, GFP-positive cell count was quantified using fluorescence microscopy and titer was determined as infectious unit per milliliter (IU ml^-1^) with the assumption that 1 GFP positive cell corresponds to 1 IU at high dilution factors given the single cycle nature of the RVP system.

### Design and cloning of CRISPR knockout library

We designed a membrane-focused CRISPR knockout library targeting genes encoding proteins predicted to be integral membrane (≥1 transmembrane segment) or GPI-anchored, using *Ae. albopictus* Foshan FPA (AalbFP1.0) gene models from VectorBase release 59 (*74*) and augmenting existing annotations with de novo predictions. Transmembrane topologies were predicted across the proteome with DeepTMHMM (*75*) v1.0.20 (release 2023-01-23) run on an HPC cluster (Harvard Medical School) inside a Singularity container with GPU passthrough (predictions were computed from the VectorBase protein FASTA and exported in GFF3 and 3-line formats; a per-protein summary is provided in Table S3). Canonical GPI-anchored proteins were identified with NetGPI 1.1 (*76*); to capture potential non-canonical cases lacking a recognizable N-terminal signal peptide, we additionally screened only the C-terminal 201 amino acids of each protein (ignoring N-terminal signal requirements) and retained sequences with predicted ω-site signatures. Beyond the membrane set, we included all genes annotated in VectorBase for GPI-anchor biogenesis, the complete kinome and phosphatome (compiled from VectorBase annotations and orthology to curated *Drosophila* sets (*77*) at https://www.flyrnai.org/tools/glad/web/), as well as smaller categories (scramblases, flippases, transferases) and a panel of benchmark controls (e.g., *Ae. albopictus* FKBP12 orthologs AALFPA_076452 and AALFPA_059142), which were included for prospective validation but not analyzed further here. Following target selection, sgRNAs were retrieved against these classes using our standard pooled-library workflow. For targets in the *Ae. albopictus* Foshan FPA assembly (AalbFP1.0), candidate guides were retrieved with CRISPR GuideXpress following our published pipeline (*19*). Briefly, all computed sgRNAs were retrieved and, where possible, up to 10 sgRNAs per gene were selected by prioritizing minimal off-target effect (OTE) score, maximal machine-learning (ML) efficiency score, and perfect concordance with the C6/36 cell-line genome (*78*) (BioProject PRJNA345486), removing any guides with mismatches to the cell-line reference. The resulting library comprises 47,677 unique sgRNAs targeting 6,361 protein-coding genes, plus 100 control sgRNAs. Oligos (109-mer) were custom synthesized on an array (Agilent), recovered by dial-out PCR using Q5 Hot Start High-Fidelity DNA Polymerase (NEB) with primers listed in Table S4. The purified amplicons were cloned by Gibson assembly (NEBuilder HiFi DNA Assembly Master Mix, NEB) into BbsI-digested pLib6.10BN-Aaeg 774 following a published protocol (*79*). The pLib6.10BN-Aaeg_774 vector was derived from pLib6.4B-Aaeg_774 by replacing the Drosophila 5C promoter with the *Ae. aegypti* ubiquitin promoter from pSL1180-HR-PUbECFP (Addgene #47917; a gift from L. Vosshall) and adjusting one BbsI site to include the first G transcribed by the native *Ae. aegypti* U6 (AAEL017774) promoter. To verify the quality of the library a two-step PCR was used to amplify the gRNA cassette and insert an inline barcode (PCR1) and Illumina sequencing adapters (PCR2) using primers specified in Table S4, Deep sequencing of the purified PCR amplicon (Illumina NovaSeq 6000, Novogene) detected 100% of designed guides (47,677/47,677 with ≥1 read), confirming successful library construction; the library is designated MFCRISPKO_AALB (Table S3) (Addgene deposition pending).

### CRISPR knockout screen

To generate the knockout pool, C6/36-HE8-Ub::Cas9-2A-Neo cells in log phase were seeded in 18 × 100-mm dishes at 3.0 × 10^7^ cells per dish in standard growth medium. After 8 h attachment, the medium was replaced and cells were transfected with a plasmid mixture containing equimolar pAeUb::ΦC31-Integrase (GenBank accession pending; Addgene pending) and the MFCRISPKO_AALB sgRNA donor library using Effectene (Qiagen; base protocol 1:50). The pAeUb::ΦC31-Integrase plasmid was generated by Gateway LR recombination of pAePUbW (Addgene #pending) with a pEntry vector bearing the ΦC31-Integrase CDS PCR-amplified from pBS130 (Addgene #26290; gift from T. Clandinin)(*80*). Across 5.4 × 10^8^ seeded cells and ∼6% recombinase-mediated cassette exchange (RMCE) recombination assessed by flow cytometry at day 25 (EBFP+), we achieved a coverage of 680 recombined cells per sgRNA (5.4 × 10^8^ × 0.06 ÷ 47,680). One day post-transfection, each dish was expanded into two 150 mm dishes in puromycin-supplemented medium and cultured for an additional 30 d with medium changes and reseeding every 4 d ; at each passage, total cell numbers were maintained above 1,000 cells per sgRNA to preserve library representation.

For screening, knockout-pool cells were plated at 4.0 × 10^7^ cells per plate (∼800-fold coverage per guide) and infected with CHIKV strain 37997 RVPs (CD20/GFP reporter) to reach 80–90% infection, as assessed by flow-cytometry for GFP. Twenty-four hours post-infection, infected cells were depleted by magnetic separation with anti-CD20 MicroBeads (Miltenyi Biotec 130-091-104). The uninfected fraction of was expanded to 4 × 10^7^ cells and reinfected for a second round to improve the signal-to-noise ratio.

Genomic DNA from the outgrown uninfected cells from each round was extracted using the QIAGEN DNeasy Blood and Tissue Kit (QIAGEN 69506) and subsequently used as template for PCR. sgRNA cassettes were amplified and sequenced on an Illumina NextSeq 1000 and gene-level enrichment analysis was conducted using MAGeCK-VISPR (version 0.5.6). Results of both rounds of the screen are provided in Table S1.

### RNAi and knockdown evaluation in mosquito cells

Unique dsRNAs (∼400–600 bp) were generated against the *Ae. aegypti* Lachesin ortholog (AAEL009295), against components of the reporter-virus particle (RVP) system RRV nsP1 and the PpluGFP2 reporter present in the replicative portion of the RVPs; positive controls for RNAi activity), and against LacZ (593 bp; bacterial β-galactosidase gene lacZ; negative control; amplified from the *Drosophila* act5C-βGal plasmid, DGRC #1220). Target regions were selected within coding sequence to avoid low-complexity/repetitive segments; Lachesin amplicons were screened to exclude off-targets using the E-RNAi web service (*81*). PCR templates were Aag2 genomic DNA for Lachesin, plasmid DNA from the RVP system, and a LacZ PCR template provided by the Drosophila RNAi Screening Center-Functional Genomics Resources (*77*) (Harvard Medical School). Amplicons were generated with 5′ T7-tailed primers (Table S4), gel-purified, and cloned into the Zero Blunt TOPO PCR vector (Thermo Fisher Scientific, 450245) for sequence verification. dsRNA was synthesized with the MEGAscript RNAi kit (Ambion AM1626) according to the manufacturer’s instructions by co-transcribing complementary strands from opposing T7 sites; products were treated with DNase, extracted with phenol:chloroform, precipitated with isopropanol, ethanol-washed, and resuspended in nuclease-free water.

For knockdown assays, Aag2 cells were seeded in standard L-15– based medium at 2.4 × 10^5^ cells per well in 12-well plates to reach ∼50% confluence the next day. Cells were transfected with 0.7 µg dsRNA per well in L-15 supplemented with 2% (v/v) FBS and no antibiotics, using Cellfectin II (Thermo Fisher Scientific, 10362100) at a 1:5 reagent:dsRNA ratio for 6 h; medium was then replaced with standard L-15 growth medium (*82*). Cells were assayed 72 h post-transfection for knockdown or subjected to RVP infection. For RVP infection, cultures were washed twice with HBSS (Gibco), and the medium was replaced with infection medium (L-15 + 5% (v/v) FBS) containing a dilution of the RVP stock to achieve an MOI of 1; GFP expression was quantified by flow cytometry 24 h post-infection.

To quantify knockdown efficiency, total RNA was extracted with TRIzol (Invitrogen), treated with DNase, and reverse transcribed from 1 µg total RNA in a 20 µl reaction using the iScript cDNA Synthesis Kit (Bio-Rad 1708890). Each cDNA was diluted 1:5 with nuclease-free water; 3 µl of diluted cDNA was used per qPCR (10 µl total) with iQ SYBR Green Supermix (Bio-Rad) under the following cycling program: 95 °C for 3 min; 39 cycles of 95 °C for 10 s and 60 °C for 30 s. Gene-specific primers for Lachesin and the housekeeping reference Rps17 (*83*) (VectorBase AAEL004175) were used at a final concentration of 300 nM and are listed in Table S4. Reactions were run in technical duplicates, specificity was verified by melt-curve analysis, and relative expression was calculated by the 2^−ΔΔCt^ method. Knockdown efficiency was reported relative to matched dsLacZ-treated controls in each biological replicate.

### Protein purification

The genes encoding *Aedes albopictus* Lachesin (residues 1–345; GenBank: XP_019932516.3), the human MXRA8 ectodomain (residues 20–337; GenBank: NP_001269511.1), the mouse MXRA8 ectodomain (residues 21-340; GenBank: NP_077225.4), the EC1 domain of human PCDH10 (residues 19–122; GenBank: NP_116586.1), the EC1 domain of sparrow PCDH10 (residues 19–122; GenBank: XP_064272571.1), and the human VLDLR ligand-binding domain (LBD) (residues 31–355; GenBank: NP_003374.3) were each cloned into the pVRC expression vector with a human IgG1 Fc tag fused at the C terminus.

To purify Lachesin–Fc, PCDH10_EC1_-Fc, and the MXRA8-Fcs, Expi293F™cells were transiently transfected with plasmids encoding the respective fusion proteins using the ExpiFectamine™293 Transfection Kit (Thermo Fisher Scientific Cat#: A14525) according to the manufacturer’s instructions. Supernatants were collected 5 d post-transfection, centrifuged at 4,000 × g for 30 min and purified using MabSelect™PrismA protein A affinity resin (Cytiva Cat#: 17549801) following the manufacturer’s protocol. The proteins were further purified by size-exclusion chromatography on a Superdex 200 increase 10/300 column (Cytiva). Proteins were stored in Tris Buffered Saline (TBS) (20 mM Tris, 150 mM NaCl, pH 7.5).

To purify VLDLRLBD-Fc, Expi293F cells were co-transfected with the pVRC vector encoding VLDLRLBD-Fc and a pCAGGS vector encoding the chaperone RAP (residues 1–353; GenBank: NP_002328). Supernatants were collected 5 d post-transfection and purified using MabSelect™PrismA protein A affinity resin as described above. To separate the VLDLR_LBD_-Fc from RAP, the column washed with 200 column volumes of 10 mM EDTA in TBS overnight. Then VLDLR_LBD_-Fc was refolded on the column by washing with 100 column volumes of 2 mM CaCl_2_ in TBS followed by elution according to the manufacturer’s protocol. The proteins were concentrated and further purified by size-exclusion chromatography on a Superdex 200 increase 10/300 GL column. RAP was stored in TBS and VLDLR_LBD_-Fc was stored in TBS containing 2 mM CaCl_2_.

The gene encoding *Aedes albopictus* lachesin (residues 1–345; GenBank: XP_019932516.3) was cloned into a pVRC vector containing a C-terminal twin-strep tag followed by an HRV 3C protease cleavage site (LEVLFQGP). Expi293F™ cells were transfected with the plasmid using the ExpiFectamine™ 293 Transfection Kit (Thermo Fisher Scientific) according to the manufacturer’s instructions. After centrifugation at 4,000 × g for 30 min, the culture supernatant was incubated with Strep-Tactin® XT Sepharose resin (IBA Lifesciences) for 1 h at 4 °C. The resin was washed with TBS, and bound proteins were eluted with 50 mM biotin in TBS. The eluted fractions were digested overnight at 4 °C with HRV 3C protease (TaKaRa, Cat. #7360). To remove the cleaved twin-strep tag and HRV 3C protease, the digested fractions were further purified by Strep-Tactin® XT Sepharose and Glutathione Sepharose™ affinity resins, respectively. The flow-through fractions were pooled, concentrated, and stored for subsequent antibody generation studies.

### Antibody generation

Anti-Lachesin antibody was generated by Biosynth International Inc. Two Specific Pathogen Free New Zealand White rabbits (12–16 weeks old) were immunized, with animal protocols approved by an internal Animal Care and Use Committee, with untagged purified *Ae. albopictus* Lachesin (residues 1–345; GenBank: XP_019932516). Five milliliters of pre-immune serum was collected prior to immunization. The immunogen (200–400 µg) was mixed at a 1:1 ratio with Freund’s adjuvants (either CFA or IFA) and administered subcutaneously per immunization. These were scheduled as follows: day 0 pre-immune bleed + boost (400 µg immunogen with CFA), day 14 boost (200 µg immunogen with IFA), day 28 boost (200 µg immunogen with IFA), day 35 production bleed #1, day 39 production bleed #2. Antibody titer of the sera from bleeds was confirmed through ELISA.

To affinity purify the polyclonal antibodies from antisera, an affinity purification column was prepared as follows. Two milligrams of untagged *Ae. albopictus* Lachesin was coupled to 2 ml AminoLink resin (Thermo Fisher Scientific 20382). Briefly, the resin was washed with phosphate buffer, then incubated with protein and NaCNBH_3_ for 6 h. The resin was washed with Tris-HCl buffer and any remaining available coupling sites were blocked by a one-hour incubation with Tris-NaCNBH_3_ buffer. The column was washed two additional times and stored at 4°C. To purify the polyclonal antibody, antisera was incubated with the prepared column for at least three hours with constant mixing followed by a wash step with salt buffer to remove non-specific material. Antibodies were eluted off the column with low pH glycine buffer, neutralized to pH 7.4. and dialyzed overnight in PBS.

### Ectopic expression construct design and generation of stable cell lines

cDNA sequences encoding human codon optimized lachesin orthologs were ordered from IDT including *Aedes albopictus* Lachesin (GenBank XP_019932516), *Aedes aegypti* Lachesin (GenBank: XP_001659911), *Culex pipens pallens* Lachesin (GenBank XP_039444183), *Culex quinquefasciatus* Lachesin (GenBank XP_001850358), *Aedes camptorhynchus* Lachesin (GenBank: XP_065093025), *Apis mellifera* Lachesin (GenBank XP_397471), *Ixodes scapularis* Lachesin (GenBank XP_029832537), *Weymonia mithii* Lachesin (GenBank: XP_055534723), *Uranotaenia lowii* Lachesin (GenBank XP_055609662), *Anopheles darlingi* Lachesin (GenBank XP_049542368), *Anopheles coustani* Lachesin (GenBank XP_058121093), and *Phlebotomus papatasi* Lachesin (GenBank XP_055707817)

cDNA encoding human MXRA8 (GenBank NM_032348), human PCDH10 (GenBank NM_032961), human VLDLR (GenBank NP_003374), mouse MXRA8 (GenBank XP_006539298), human LDLRAD3 (GenBank AAI43825), human NEGR1 (GenBank NP_776169), and human LSAMP (GenBank KAI2530963) were ordered from IDT.

*Passer domesticus* PCDH10 and MXRA8 sequences were obtained by aligning the coding sequences of *Passer montanus* PCDH10 (GenBank XM_039733439.1) or MXRA8 (GenBank XM_039727729.1) against the genome of *P. domesticus* (GenBank GCA_001700915.1) and assembling aligned fragments.

N-terminal Flag constructs for domain mapping were constructed using the codon optimized *Ae*.*albopictus* lachesin as above (GenBank: XP_019932516) with the Flag tag (DYKDDDK) inserted after the signal peptide (between A22 and Q23) The remaining constructs all contained the signal peptide and Flag tag as above with the following inclusions: LacΔD3-Flag construct contained (Q23–F224 + I331–F366), LacΔD1-Flag (P142–V330 + I331–F366), D1-Flag (Q23–V131 + I331–F366), D2-Flag (P142–F232 + I331–F366), and stalk-Flag (I331–F366).

The above constructs were cloned into lentiGuide-Puro (Addgene #52963) and transfected with psPAX2 (Addgene #12260) and PMD2.G (Addgene #12259) at a ratio of 3:2:1 into HEK 293T cells using Lipofectamine 3000 (Thermo Fisher Scientific). Lentiviruses were collected 2 d post-transfection and used to transduce K562 cells. Transduced K562 cells were selected using puromycin at 2 µg ml^-1^. Cell lines were confirmed to express the transduced constructs by cell surface immunostaining.

### Cell surface antibody staining

Cells were blocked in blocking buffer (5% (v/v) goat serum in PBS) for 30 min at 4°C and washed once with binding buffer (2% (v/v) goat serum in PBS). Primary antibodies were diluted to 5 µg ml^-1^ in binding buffer. Primary antibodies used include polyclonal anti-PCDH10 (Proteintech 21859-1-AP), anti-VLDLR (GeneTex GTX79552), anti-MXRA8 (MBL International W040-3), anti-NEGR1 (Invitrogen PA5-52870), anti-LDLRAD3 (Invitrogen BS-18209R), anti-Rabbit IgG polyclonal isotype (proteintech 30000-0-AP), anti-SFV immune ascitic fluid (ATCC VR-1247AF) or control ascitic fluid (ATCC VR-1247CAF). Cells were washed twice in wash buffer and were subsequently incubated with PE-conjugated donkey anti-rabbit F(ab′)2 fragment (Jackson ImmunoResearch 711-116-152) or a PE-conjugated donkey anti-mouse F(ab′)2 fragment (Jackson ImmunoResearch 715-116-150) diluted 1:200 in binding buffer for 30 min at 4 °C. Cells were then washed twice in wash buffer, followed by two washes in PBS before being fixed in 1% (v/v) formalin in PBS. Cell surface receptor expression was detected using an iQue3 Screener PLUS (Intellicyt) with ForeCyt (Sartorius) software (version 8.1.7524). Antibody staining was visualized using FlowJo (version 10.6.2).

For cells expressing Flag tagged constructs, a commercial APC-conjugated anti-Flag (DYKDDDK) antibody (Biolegend 637308) or isotype control (Biolegend 402306) was diluted to 5 µg ml^-1^ in binding buffer as above after a 30 min block in blocking buffer. Cells were then washed twice in binding buffer followed by two washes in PBS before being fixed in 1% (v/v) formalin in PBS. Cell surface Flag staining was detected using an iQue3 Screener PLUS (Intellicyt) with ForeCyt (Sartorius) software. Antibody staining was visualized using FlowJo (version 10.6.2).

For cells transfected with alphavirus glycoproteins, HEK 293T cells were transfected with the E3–E2–(6K/TF)–E1 sequences of the indicated alphaviruses in the pCAAGS or pTWIST-CMV-BG-WPRE-Neo expression vectors from the “reporter virus particle generation” Methods section above. Two days post transfection, cells were harvested and blocked in blocking buffer for 30 min at 4 °C and washed once with binding buffer (2% (v/v) goat serum in PBS). Diluted Fc fusion proteins were added at the indicated concentrations and incubated for 30 min at 4°C before cells were washed twice with washing buffer. Cells were subsequently incubated with PE-conjugated donkey anti-human F(ab′)2 fragment (Jackson ImmunoResearch 109-116-098) diluted 1:200 in binding buffer for 30 min at 4 °C. Cells were washed twice in wash buffer, followed by two washes in PBS before being fixed in 1% (v/v) formalin in PBS. Fc fusion protein staining was detected using an iQue3 Screener PLUS (Intellicyt) with ForeCyt (Sartorius) software. Antibody staining was visualized using FlowJo (version 10.6.2). Alphavirus E3–E2–(6K/TF)–E1 sequences used only for staining were cloned into the pTWIST-CMV-BG-WPRE-Neo expression vector and are UNAV strain BeAr13136 (GenBank AF339481), and BEBV strain MM2354 (GenBank AF339480).

### Inhibition of RVP entry using recombinant proteins

Fc fusion recombinant proteins were serially diluted at half-logs and incubated with RVP before being placed on cells at MOI 1. At 24 h post infection, cells were washed twice with PBS and fixed in 1% formalin in PBS (v/v). GFP positivity was used as a proxy for RVP entry and was quantified using an iQue3 Screener PLUS (Sartorius) running the IntelliCyt ForeCyt Standard Edition (Sartorius) software (version 8.1.7524). Relative infection was calculated as: % Relative Infection = (%GFP positive cells in the presence of protein)/(%GFP positive cells in the absence of protein) × 100.

### Single-cycle recombinant VSV generation

Single-cycle VSV was rescued from pBS-fT plasmids encoding VSV-N (pBS-N), VSV-P (pBS-P), VSV-L (pBS-L), and VSV-G (pBS-G) and a plasmid encoding the antigenome of VSV with G replaced with GFP (pVSV-ΔG-GFP). All expression vectors contain a T7 promoter (Kerafast EH1004). BHK21/WI2 cells (Kerafast EH1011) were seeded in a 6-well plate and infected with vTF7-3 vaccinia virus (VACV) encoding T7 polymerase for 1 h with rocking every 15 min. After 1 h, VACV containing media was removed, 1 ml of fresh media was added per well and cells were transfected in an 8:3:5:1:8 ratio of pVSV-ΔG-GFP, pBS-N, pBS-P, pBS-L, and pBS-G using Lipofectamine 3000 according to manufacturer’s instructions (Thermo Fisher Scientific). Two days post-transfection, pVSVΔG was collected from culture supernatant and filtered through 0.22 µm filter to remove residual vTF7-3 VACV.

To amplify rescued virus, BHK21/WI2 cells were seeded in a 6-well plate and transfected with 2 µg PMD2.G (Addgene #12259) the following day. The next day, the media on transfected cells was changed to 1 ml of FBS-free DMEM supplemented with HEPES and cells were infected 1 ml of pVSVΔG for 1 h with rocking every 15 min at 37 °C. After 1 h, the inoculum was removed, and cells were washed twice with DMEM + 10% (v/v) FBS + 25 mM HEPES. 1.5 ml of DMEM + 10% (v/v) FBS + 25 mM HEPES was added to each well. The following day, culture supernatant was collected from each well and filtered through 0.22 µm filter to remove potential residual vTF7-3 VACV, aliquoted, and stored at -80 °C

### Phospholipase C treatment and RVP infection

C6/36-HE8C cells were seeded at 3 × 10^5^ cells per well in a V-bottom 96-well plate. Cells were washed twice with PBS and 150 µl of phospholipase C (Invitrogen) diluted to 2 U ml^-1^ in PBS was added to each well. Cells were incubated for 1 h at 37°C. Cells were then washed twice with PBS before being infected with RVP or VSVDG-G inoculum (MOI=10 as titered on C6/36-HE8C cells) for 1 h at 4 °C with rocking. After 1 h, the inoculum was removed, and cells were washed four times in PBS before adding 100 µl culture media and being plated in a 96-well plate. Infection was quantified by measuring GFP 24 h post-infection using flow cytometry using an iQue3 Screener PLUS (Intellicyt) with ForeCyt (Sartorius) software. Data were visualized using ForteBio Data Analysis HT version 12.0.1.55 software.

### Biolayer interferometry

Biolayer interferometry was performed using an Octet RED96e (Sartorius), and data were analyzed with ForteBio Data Analysis HT software. Lachesin–Fc and *Hs*PCDH10_EC1_–Fc were immobilized onto Anti-Human IgG Fc Capture (AHC) Biosensors (Sartorius 18-5063) at a concentration of 250 nM in kinetic buffer (TBS containing 0.1% (w/v) BSA and 0.01% (v/v) Tween) for 300 s. The sensor tips were then dipped into the kinetic buffer for a baseline measurement of 60 s, followed by immersion in a 1 µM solution of CHIKV or SFV VLPs for 300 s, and finally transferred back to the kinetic buffer for a 300 s dissociation phase.

### Virus-like particle generation

We transfected plasmids encoding the structural polyprotein (capsid–E3– E2–(6K/TF)–E1) of CHIKV strain 37997 (*84*) (GenBank: AAU43881), WEEV strain CBA87 (GenBank: DQ432026), which contains a nuclear-localization mutation in the gene encoding the capsid protein to increase expression (*85*), into Expi293F™ cells using the ExpiFectamine 293 Transfection Kit (Thermo Fisher Scientific) according to the manufacturer’s protocol. Culture supernatant was collected 5 d post-transfection and clarified of cell debris by centrifugation at 3,000 × *g* for 20 min. The clarified supernatant was ultracentrifuged with a sucrose cushion consisting of 5 ml of 35% (w/v) sucrose and 5 ml of 70% (w/v) sucrose at 150,000 × *g* for 3 h in a Beckman SW32Ti rotor at 4 °C. The VLPs were pooled from the interface of the 35% (w/v) and 70% (w/v) sucrose cushions. The VLPs were then loaded onto a 20%–70% (w/v) continuous sucrose density gradient and ultracentrifuged for 1.5 h in a Beckman SW41 rotor at 210,000 × *g* at 4 °C.

We produced SFV VLPs by transiently transfecting HEK 293T cells grown in adherent culture with a vector encoding the structural polyprotein of SFV strain SFV4 (GenBank: AKC01668.1) using lipofectamine 3000 (Invitrogen) according to the manufacturer’s instructions. Culture supernatant was collected 48 h post-transfection. The cell debris were removed by centrifugation for 20 min at 3,000 × *g*. The VLP was centrifuged with a 30% (w/v) sucrose cushion at 110,000 × *g* for 2.5 h in a SW32Ti rotor (Beckman) at 4 °C. The VLP pellet was resuspended in DPBS (Thermo Fisher Scientific Cat#: 14190-144) and purified by ultracentrifugation on a 20–70% (w/v) sucrose density gradient in a Beckman SW41 rotor at 210,000 × *g* for 1.5 h at 4 °C. The VLP band was collected from the gradient and stored at 4 °C without buffer exchange and not frozen. We confirmed particle integrity and the absence of degradation products using SDS–PAGE. VLPs were always used within seven days of purification, and buffer exchanged based on the application immediately before use.

### VLP fluorescent labeling

After purification, VLPs were buffer exchanged into 0.1 M Sodium bicarbonate pH 8.3 and diluted to a concentration of 1 mg ml^-1^. Prior to VLP labeling, Alexa Fluor 647 (AF647) NHS ester (Invitrogen A37573) was resuspended to a concentration of 1 mg ml^-1^ in dimethyl sulfoxide (DMSO). Twenty-five micrograms of AF647 was added to 1 mg of VLPs and incubated at room temperature for 30 min. To remove excess AF647 from the solution after labeling, the solution was flowed through a Zeba Spin Desalting Column (Thermo Fisher Scientific 89877) and VLPs were buffer exchanged into PBS. Labeled VLPs were stored at 4 °C until used for confocal microscopy. To confirm integrity of the VLPs after labeling, we performed SDS-PAGE on these VLP samples in reducing and non-reducing conditions and imaged the bands in the far-red range. Only glycoproteins E1 and E2, not capsid, were labeled with 647, indicating proper labeling and intact VLPs (Fig. S9A). Labeled VLPs were used within 6 h of labeling.

### Confocal microscopy

A total of 10^7^ K562 cells stably expressing either human MXRA8, human PCDH10, or *Ae. Albopictus* Lachesin were centrifuged at 300 × g for 5 min and resuspended in a heparinase mixture [heparinase I (Sigma-Aldrich H2519) at 4 units ml^-1^, heparinase II (Sigma-Aldrich H6512) at 2 units ml^-1^, and heparinase III (Sigma-Aldrich H8891) at 4 units ml^-1^]. Cells were treated with heparinases for 1 h at 37 °C. Cells were then washed and resuspended at a density of 0.5 × 10^6^ ml^-1^ in RPMI1640 supplemented with 2% (v/v) FBS and 25 mM HEPES. Twenty-five micrograms of labeled VLP were added to 0.5 × 10^6^ cells. For cells kept a 4 °C, cells were kept on ice after the addition of VLPs. For cells kept at 37 °C, cells were incubated in a heat block at 37 °C. Cells were incubated with VLPs for 30 min before being washed once with PBS. Immediately before imaging, cells were treated with 500 µl Alexa Fluor 488-conjugated wheat germ agglutinin (WGA-AF488) (Invitrogen W11261) at 1 µg ml^-1^ in PBS, washed once with cold PBS before being resuspended in 80 µl of cold PBS. Cells were then plated in glass bottom microwell dishes (MatTek P35G-1.5-14-C) and imaged within 5 min of plating.

Cells were imaged with a Yokogawa CSU-W1 single disk (50 µm pinhole size) spinning disk confocal unit attached to a Nikon Ti2 inverted microscope equipped with a Nikon linear-encoded motorized stage, an Andor Zyla 4.2 plus (6.5 µm photodiode size) sCMOS camera using a Plan Apo 60x/1.3 silicon immersion with the correction collar set to minimize spherical aberration. The final digital resolution of the image was 0.1 µm per pixel. Fluorescence from WGA-AF488 and VLPs conjugated to AF647 was collected by illuminating the sample with a solid state directly modulated 488 nm diode 100 mW (at the fiber tip) laser line or a solid state, directly modulated 640 nm diode 100 mW (laser tip) laser line in a Nikon LUNF XL laser combiner, respectively. A hard-coated Semrock Di01-T405/488/568/647 multi-bandpass dichroic mirror was used for both channels. Signal from each channel was acquired sequentially with either a hard-coated Chroma ET525/36 m or Chroma ET700/75 m emission filters in a filter wheel placed within the scan unit, respectively. z-stacks were set by determining the top and bottom of the cell, using WGA-AF488 fluorescence as a reference. The approximate z-distance was 15 µm, and the step size was set to 0.2, using the piezo stage insert (Mad City Labs, 500 µm). Fluorescence from each fluorophore was acquired sequentially at each z-step of the confocal to improve the axial precision of the measurements. Nikon NIS-Elements Advanced Research (AR) version 5.02 acquisition software was used to acquire the data, and the files were exported in ND2 file format. Figures were generated using Fiji (*86*). Max intensity projection (MIP) renderings were created by using the 3D projection function (Stacks>3D Project) with 10° increments and interpolation selected to smooth the 3D rendering

### Infection of stable cell lines with infectious reporter alphaviruses

K562 cells transduced with an empty lentiGuide-Puro vector or transduced to overexpress human MXRA8, PCDH10, VLDLR, LDLRAD3, or *Ae. albopictus* Lachesin were seeded onto poly-L-lysine coated 96-well plates (1 ×10^5^ per well) and allowed to settle for 3 h. The medium was replaced with 100 µl of a solution containing 1 × 10^5^ FFU infectious reporter-alphaviruses in infection medium (RPMI-1640 supplemented with 3% (v/v) FBS, 25 mM HEPES, 1% (v/v) penicillin-streptomycin) at an MOI of 1 (with exception of CHIKV WA, inoculated at an MOI of 0.5). After 2 h of inoculation at 37 °C with 5% CO2, the cells were washed once with 100 µl DPBS, followed by the addition of 100 µl infection medium. Fluorescent reporter expression was followed for an additional 24 h using the Incucyte S3 imaging platform (Sartorius) in a humidified incubator at 37 °C with 5% CO_2_, using 10× magnification and acquisition of 4 images per well every 4 h. Data were analyzed using the manufacturer’s Basic Analyzer tool and exported as integrated fluorescent intensity (in Green or Red Calibrated Units) per image normalized for phase confluence. At 26 hours post infection, the cells were washed with 100 µl DPBS and detached from the plate by forceful resuspension. Technical replicates were pooled. Cell suspensions were fixed in 4% PFA for 30min at RT and resuspended in DPBS with 2% (v/v) FCS and 2 mM EDTA for determination of reporter-positive cell percentages by flow cytometry on an ID7000 5-laser Spectral Cell Analyzer (SONY), and analyzed using FlowJo v10.10.0 (BD Life Sciences).

### Replication kinetics of SFV A774wt in stable cell lines

K562 cells transduced with an empty lentiGuide-Puro vector or transduced to overexpress human MXRA8, PCDH10, VLDLR, or *Ae. albopictus* Lachesin were infected with SFV A774wt at an MOI of 0.01 in maintenance media [RPMI with 2% (v/v) FBS, 25mM HEPES, and 1% (v/v) penicillin-streptomycin]. After one hour incubation at 37 °C with 5% CO_2_, the inoculum was removed, and cells were washed twice with DPBS prior to a final resuspension in maintenance media. Samples were collected immediately following this resuspension, as well as at 12, 24, and 48 h post-infection with the harvested volume replaced by fresh maintenance media. Samples were stored at -80 °C until titration via plaque assay on Vero cells.

### Generation of phylogenetic trees

For the tree of CHIKV lineages, sequences encoding the E2–(6K/TF)–E1 proteins of 42 CHIKV strains with full genome sequences available (Table S2) were aligned in MEGA12 using the built-in MUSCLE algorithm. A maximum-likelihood phylogenetic tree was constructed using the aligned sequences with the LG model (+freq)(*87*). The evolutionary rate differences among sites were modeled using a discrete Gamma distribution across 4 categories (*+G*, parameter = 2.3703) The bootstrap method was used to test phylogeny with 1000 bootstrap replications. For a list of CHIKV E2-6K-E1 sequences used in the generation of the phylogenetic tree, see Table S4.

For the tree of Lachesin orthologs, full-length amino acid sequences of 13 Lachesin orthologs were aligned in MEGA11 using the built-in MUSCLE algorithm. A maximum-likelihood phylogenetic tree was constructed for the aligned sequences using the LG model with discrete Gamma distribution across 4 categories. The bootstrap method was used to test phylogeny with 1000 bootstrap replications.

For the tree of alphaviruses, amino acid sequences of the E2– (6K/TF)–E1 proteins of 15 alphaviruses were aligned in MEGA11 using the built-in MUSCLE algorithm. A maximum-likelihood phylogenetic tree was constructed for the aligned sequences using the LG model with discrete Gamma distribution across 4 categories. The bootstrap method was used to test phylogeny with 1000 bootstrap replications.

---

**Figure S1.**
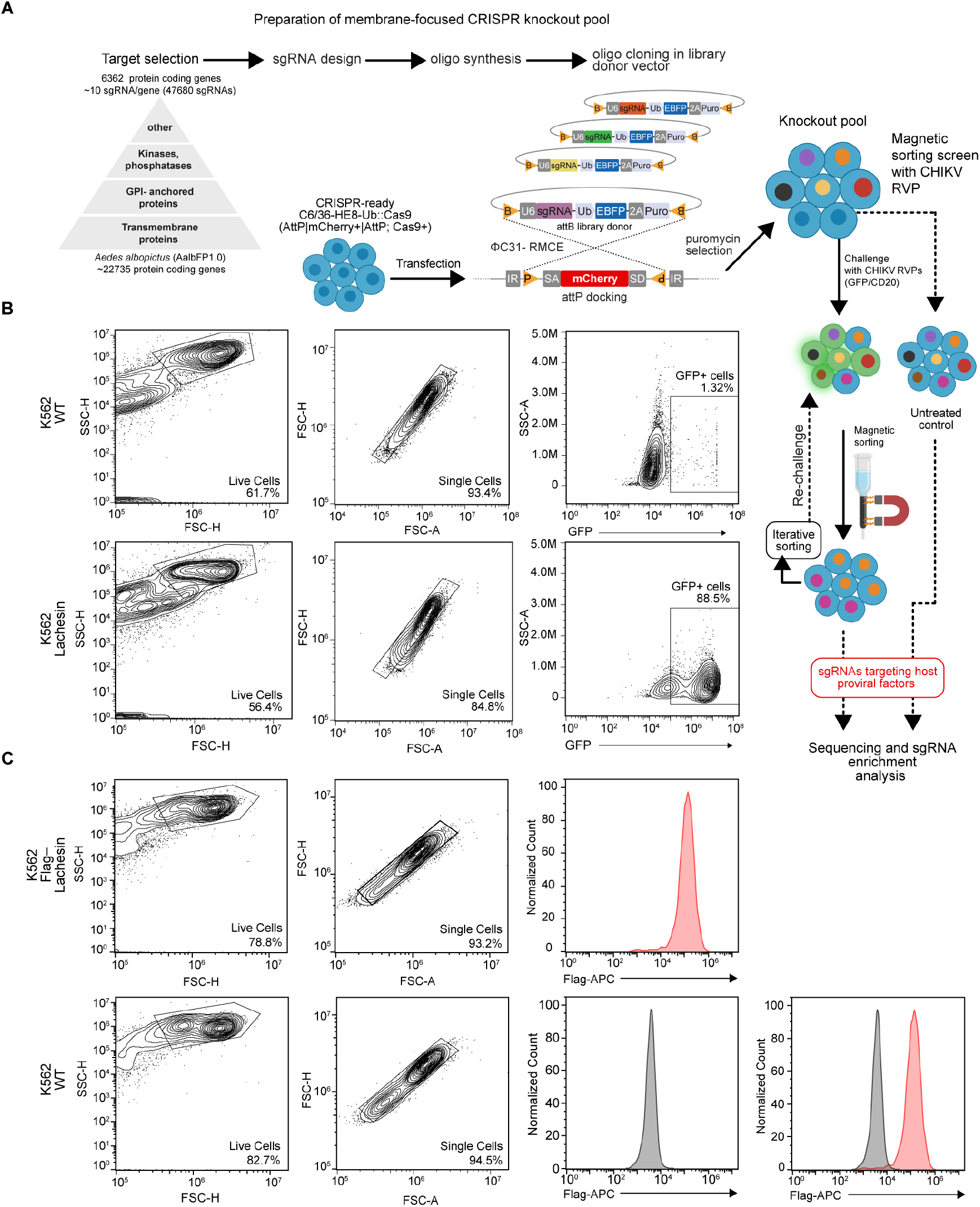
CRISPR-Cas9 screening workflow and flow cytometry gating. **(A)** Schematic diagram of the membrane-focused CRISPR-Cas9 knockout screen in *Ae. albopictus* C6/36-HE8-Ub::Cas9-2A-Neo cells. A pooled library (47,680 sgRNAs targeting 6,362 genes) was designed with CRISPR GuideXpress, cloned into an attB donor library vector, and delivered by ΦC31-mediated recombinase-mediated cassette exchange (RMCE). After approximately 30 d of puromycin selection, the knockout pool (∼800× coverage per sgRNA) was challenged with CHIKV 37997 reporter viral particles (RVPs) encoding CD20–GFP. Twenty-four hours post-infection, infected cells (CD20^+^) were removed using anti-CD20 magnetic beads; the uninfected fraction (CD20^−^) was expanded and rechallenged once. Genomic DNA from the CD20 depleted fractions and from an uninfected knockout-pool input control was used to PCR-amplify sgRNA cassettes for sequencing, and sgRNA enrichment was analyzed at the gene level with MAGeCK-VISPR (*1*). **(B)** Example of the flow cytometry gating strategy for quantification of GFP-expression in cells infected with GFP-expressing CHIKV (strain 37997) RVPs. Wild-type (WT) K562 cells (top panels) or K562 cells overexpressing *Ae. albopictus* Lachesin (bottom panels) are shown. **(C)** Flow cytometry gating strategy for monitoring of cell surface expression of alphavirus receptors. K562 cells expressing *Ae. albopictus* Lachesin-Flag (top panels) or WT K562 cells (bottom panels) were stained with an anti-Flag-APC-conjugated antibody for staining and detection. Right-most panel is an overlay of WT and Lachesin-Flag K562 cells.

**Figure S2.**
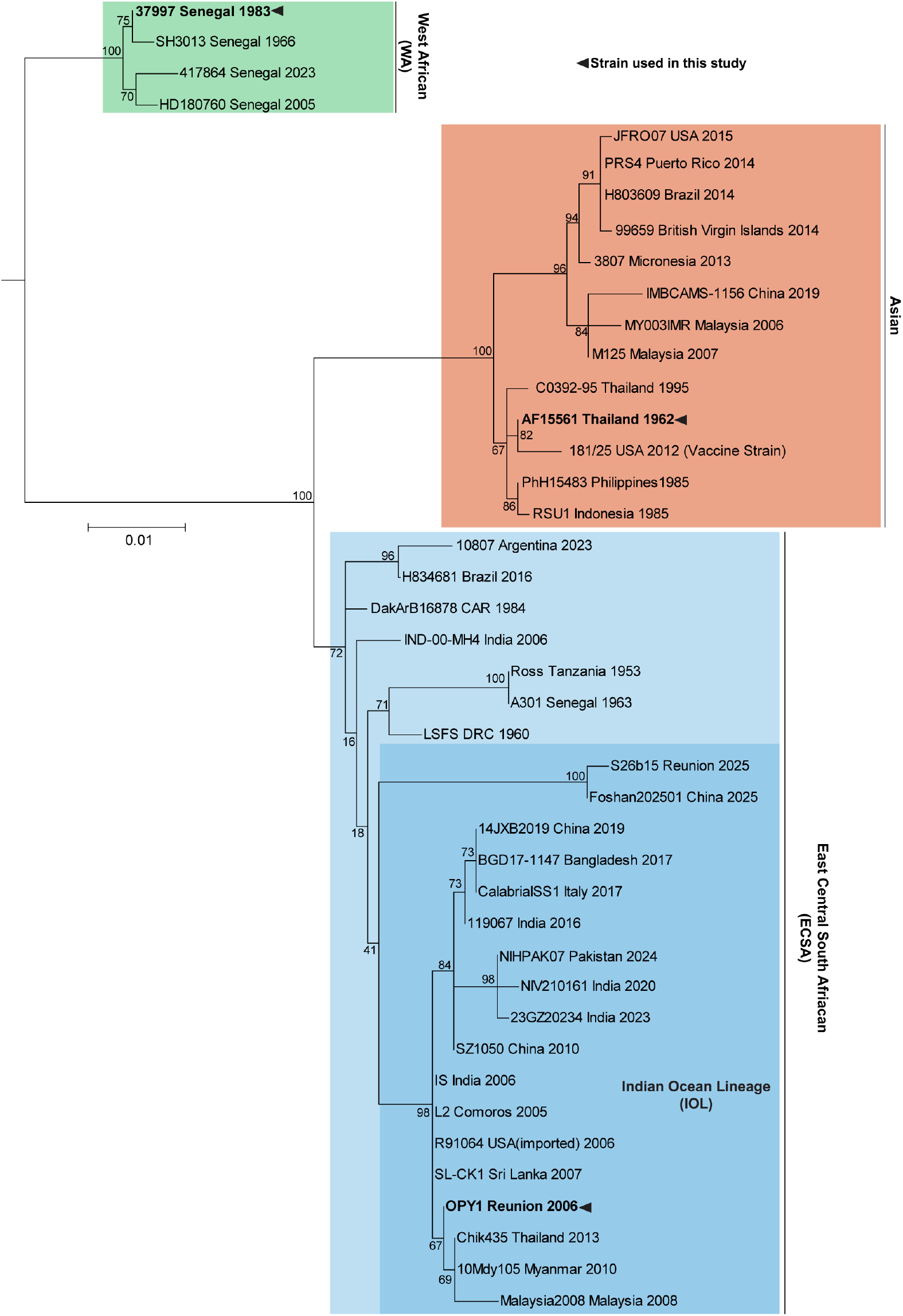
Phylogenetic analysis of CHIKV strains. Maximum likelihood phylogenetic tree of 42 CHIKV strains with full genome sequences available, using the coding sequences of the structural polyprotein genes (Table S2). The tree is rooted with WEEV strain EQ1090 structural polyprotein genes (GenBank: WYA89767) and ONNV strain UVRI0804 (Genbank: UUB88401) as outgroups (not shown). Numbers at nodes indicate bootstrap values. In cases in which the branches are too small bootstrap values may not be shown. Scale bar represents 0.01 amino acid substitutions per site. Taxon labels include strain name, country of isolation, and year of isolation. Different clades are colored and labeled. Strains we examined in this study are bolded and indicated by a triangle.

**Figure S3.**
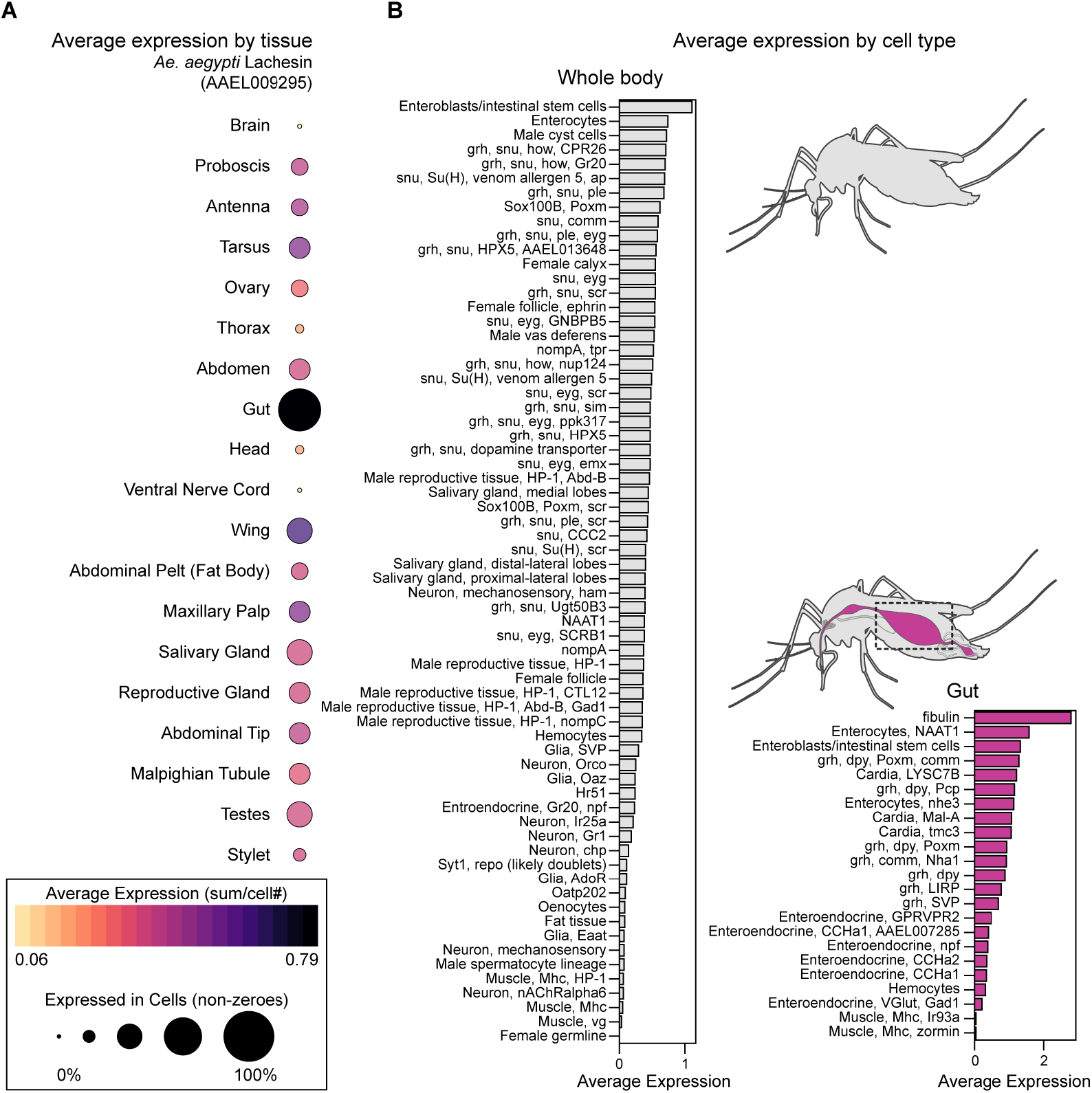
Expression of *Ae. aegypti* Lachesin across tissues and cell types. (**A)** Average expression of *Ae. aegypti* Lachesin (Vectorbase: AAEL009295) across adult tissues, visualized as dot plots from the *Ae. aegypti* single-cell atlas using the UCSC Cell Browsers (*88*). The sizes of the circles represent the percentage of cells from each tissue that expresses Lachesin. The colors of the circles represent the average expression of Lachesin across all cells from a given tissue. **(B)** Average expression of *Ae. aegypti* Lachesin across all annotated cell types in the whole body (left) and within gut cell populations (right). Data were extracted from Goldman et al. (BioProject accession PRJNA1223381) (*27*). Expression data were visualized using the UCSC Cell Browser and summarized as bar plots showing average normalized expression per tissue and per cell type. Mosquito illustrations were generated with BioRender.com.

**Figure S4.**
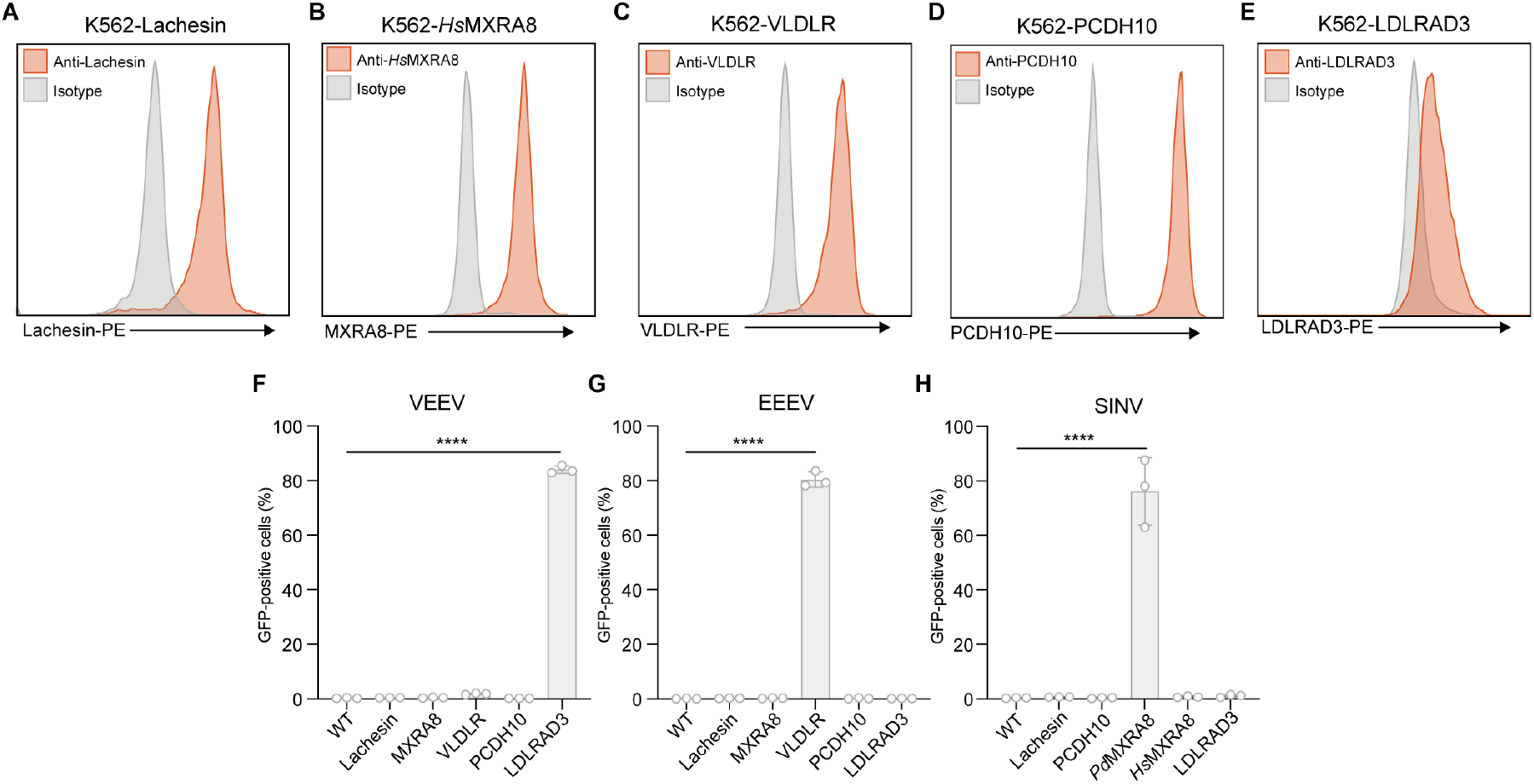
Alphavirus receptor staining and encephalitic alphavirus RVP infections. **(A–E)** Immunostaining of K562 cells stably expressing *Ae. albopictus* Lachesin (**A**), human MXRA8 (**B**), human VLDLR (**C**), human PCDH10 (**D**), or human LDLRAD3 (**E**). **(F,G)** Wild-type (WT) K562 cells or K562 cells stably expressing *Ae. albopictus* Lachesin, human MXRA8, VLDLR, PCDH10, or LDLRAD3 were infected with VEEV (INH-9813) (**f**) or EEEV (FL91-469) GFP-expressing RVPs and infection was measured using flow cytometry 24 h post-infection. **(H)**, WT K562 or K562 cells expressing *Ae. albopictus* Lachesin, human PCDH10, avian (*Passer domesticus*) MXRA8, human MXRA8, or human LDLRAD3 were infected with GFP-expressing SINV (TR339) RVPs and infection was measured 24 h post-infection using flow cytometry.Data are mean ± s.d. of 3 independent experiments performed in duplicate (*n* = 3) (**F**–**H**). One-way ANOVA Dunnett’s multiple comparison \*\*\*\**P* < 0.0001 (**F**–**H**)

**Figure S5.**
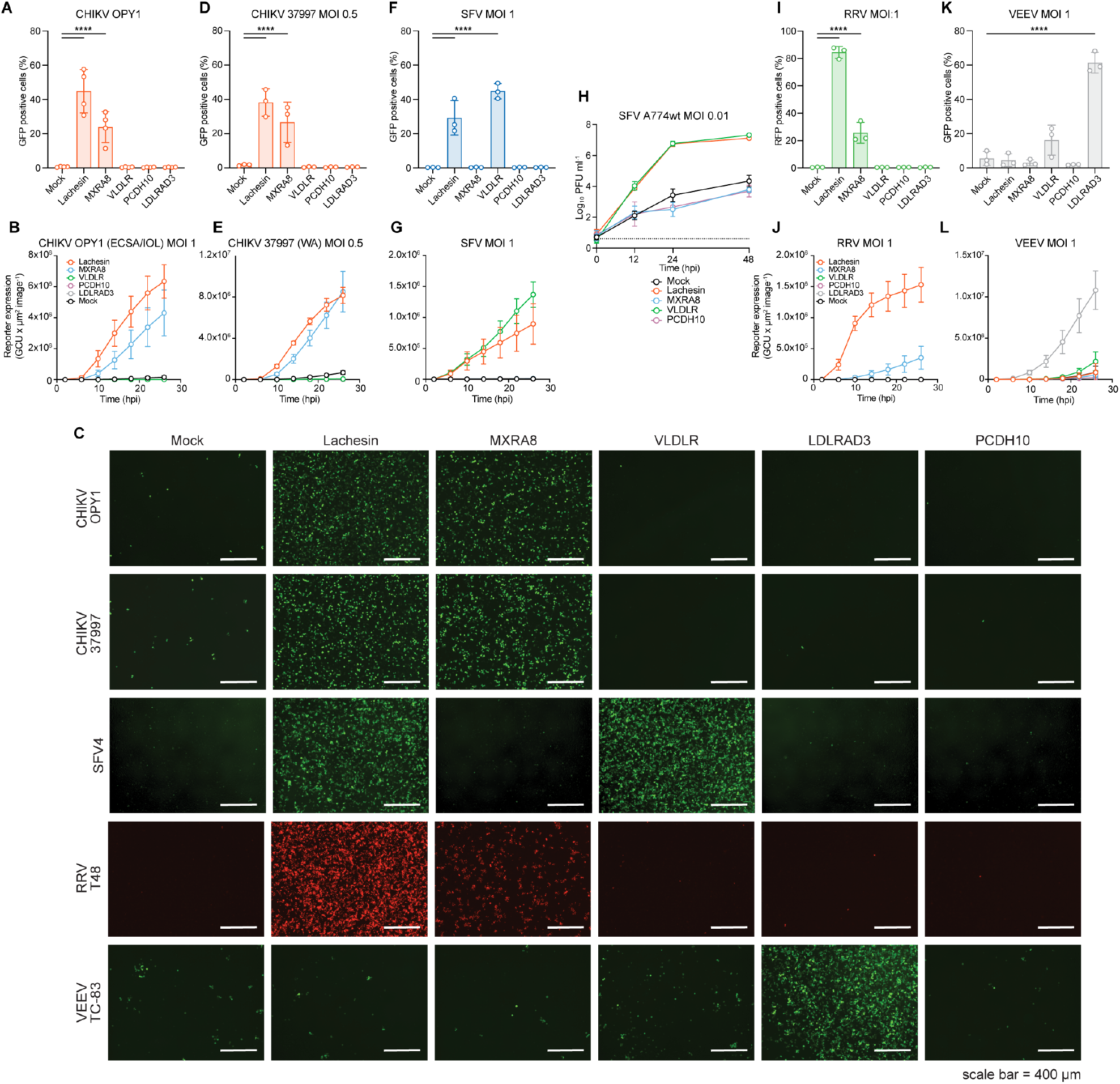
Lachesin facilitates infection by replication-competent alphaviruses. **(A,B)** Mock-transduced K562 cells or K562 cells expressing *Ae. albopictus* Lachesin or the indicated human receptors were infected with replication-competent CHIKV OPY1 expressing GFP and infection was monitored using flow cytometry 26 h post-infection (**A**) or by using a live-imaging system to monitor reporter expression (**B**). **(C)** Representative images of cells from (**B, E, G, J**, and **l**), taken with a live-cell imaging system 26 h post infection from the experiments. **(D,E)** Mock-transduced K562 cells or K562 cells expressing *Ae. albopictus* Lachesin or the indicated human receptors were infected with replication-competent CHIKV strain 37997 expressing GFP. Infection was monitored using flow cytometry 26 h post-infection (**D**) or by using a live-cell imaging system to monitor reporter expression (**E**). **(F,G)** Mock-transduced K562 cells or K562 cells expressing *Ae. albopictus* Lachesin, or the indicated human receptors were infected with replication-competent SFV strain SFV4 expressing GFP. Infection was monitored using flow cytometry 26 h post-infection (**F**) or by using a live-cell imaging system to monitor reporter expression (**G**). **(H)** Viral replication curve for replication-competent SFV strain A774wt in mock-transduced K562 cells or K562 cells expressing *Ae. albopictus* Lachesin or the indicated human receptors. **(I,J)** Mock-transduced K562 cells or K562 cells expressing *Ae. albopictus* Lachesin or human receptors were infected with RRV strain T48 expressing red fluorescent protein (RFP). Infection was monitored using flow cytometry 26 h post-infection (**I**) or by using a live-cell imaging system to monitor reporter expression at different time points (**J**). **(K,L)** Mock-transduced K562 cells or K562 cells expressing *Ae. albopictus* Lachesin or human receptors were infected with VEEV strain TC-83 expressing GF. Infection was monitored using flow cytometry 26 h post-infection (**K**) or by using a live-cell imaging system to monitor reporter expression at different time points (**L**). In **B, E, G, J**, and **L**, reporter expression was monitored over time as integrated fluorescent intensity (in Green or Red Calibrated Units) per image normalized for phase confluence as assessed using an Incucyte (Sartorius) live-cell imaging system. Data are mean ± s.d. of three independent experiments performed in triplicate (*n* = 3), with the exception of CHIKV-OPY1, for which data are mean ± s.d. of 4 independent experiments performed in triplicate for (*n* = 4). One-way ANOVA with Dunnett’s multiple comparison test. *\*\*\*\*P <* 0.0001 (**A, D, F, I**, and **K**).

**Figure S6.**
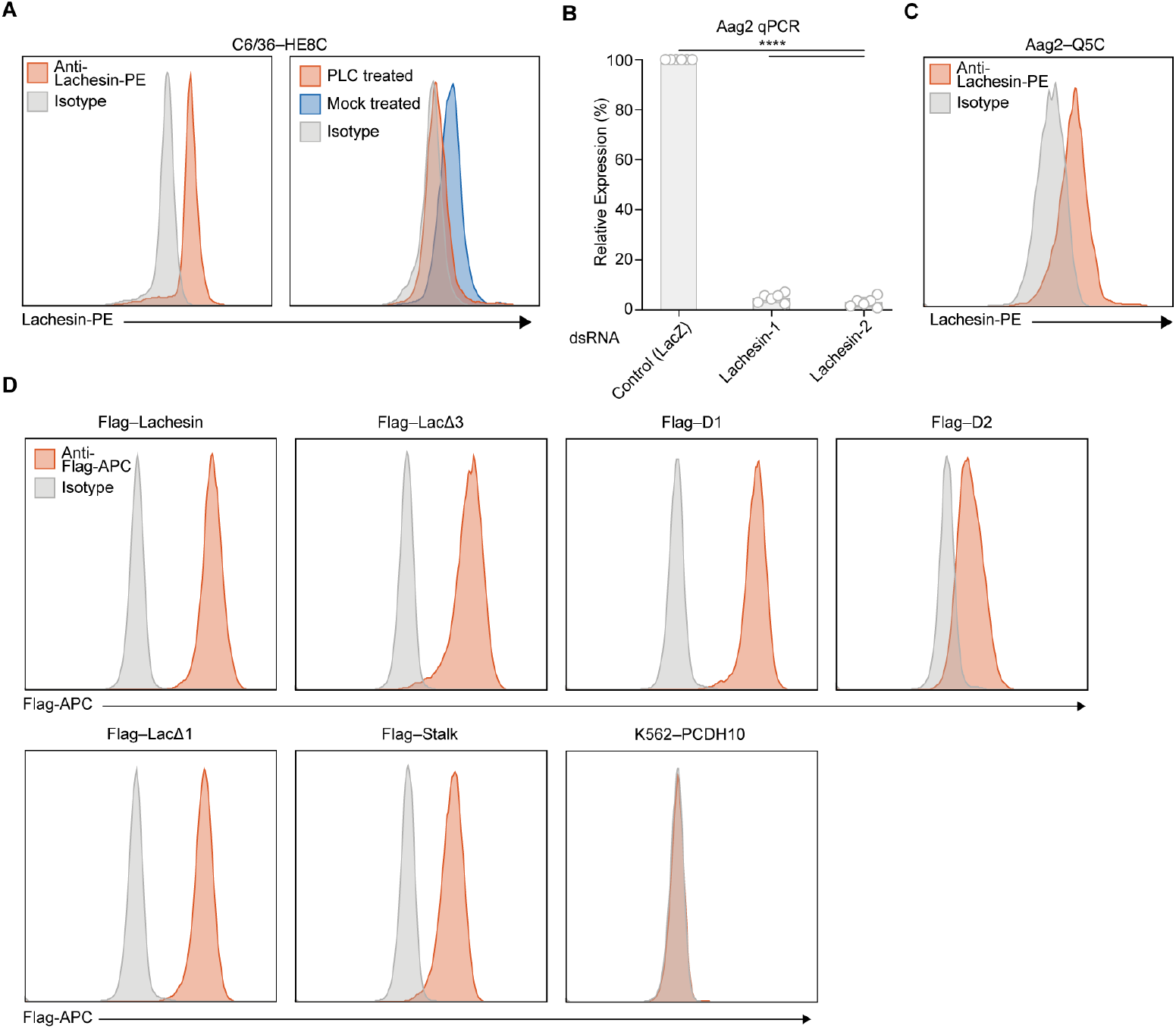
Lachesin and N-terminal Flag domain construct cell surface staining and PLC and dsRNA knockdown validation. **(A)** Lachesin cell surface immunostaining of C6/36-HE8C cells (left panel) or mock treated or phospholipase C (PLC)-treated C6/36-HE8C cells (right panel). The experiment was performed three times and representative data are shown. PE: *R*-phycoerythrin. **(B)**, Expression of Lachesin in Aag2 cells treated with dsRNA targeting Lachesin. Aag2 cells were transfected with dsRNAs targeting Lachesin (Lachesin-1, Lachesin-2) or dsRNA targeting β-galactosidase (LacZ). RNA was extracted and reverse transcribed and subjected to qPCR analysis. The experiment was performed six times and normalized to Lachesin expression of cells treated with dsRNA targeting LacZ. **(C)** Cell surface immunostaining of *Ae. aegypti* Aag2 cells with *Ae. albopictus* Lachesin polyclonal antibody. **(D)** Cell surface immunostaining of K562 cells stably expressing Lachesin-Flag, LacΔD3-Flag, LacΔD1-Flag, D1-Flag, D2-Flag, Stalk-Flag or PCDH10 (control). Data are mean ± s.d. of six experiments performed in triplicate (*n* = 6); one-way ANOVA with Dunnett’s multiple comparisons test, *\*\*\*\*P* < 0.0001.

**Figure S7.**
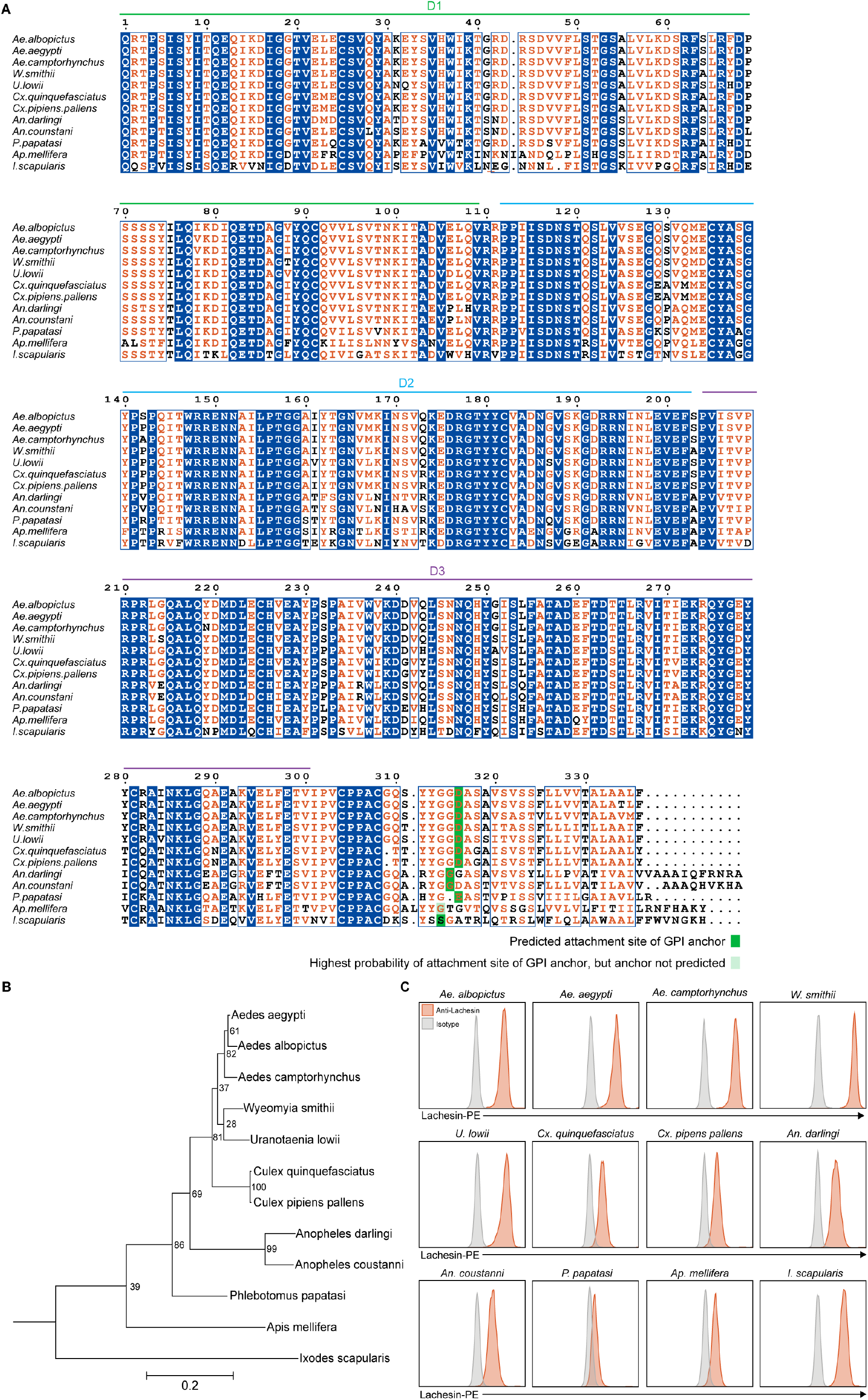
Multiple sequence alignment, phylogenetic tree, and cell staining of Lachesin orthologs. **(A)**, Multiple sequence alignment of Lachesin orthologs used in this study, generated with Espript 3.0 (*89*). GPI anchoring sites were predicted using NetGPI 1.1(*76*). Residues that are completely conserved in all aligned sequences have a blue background. Boxed residues show positions where a single majority residue or multiple chemically similar residues are found. Such residues are in red.**(B)** Maximum likelihood phylogenetic tree of Lachesin orthologs used in this study, using the Lachesin CDS. The tree is rooted with *Paramacrobiotus metroplianus* (Tartigrade) (not shown) Lachesin as an outgroup. Numbers at nodes indicate bootstrap values. Scale bar represents 0.2 amino acid substitutions per site. **(C)** Cell surface immunostaining of K562 cells stably expressing the indicated Lachesin orthologs with anti-Lachesin polyclonal antibody.

**Figure S8.**
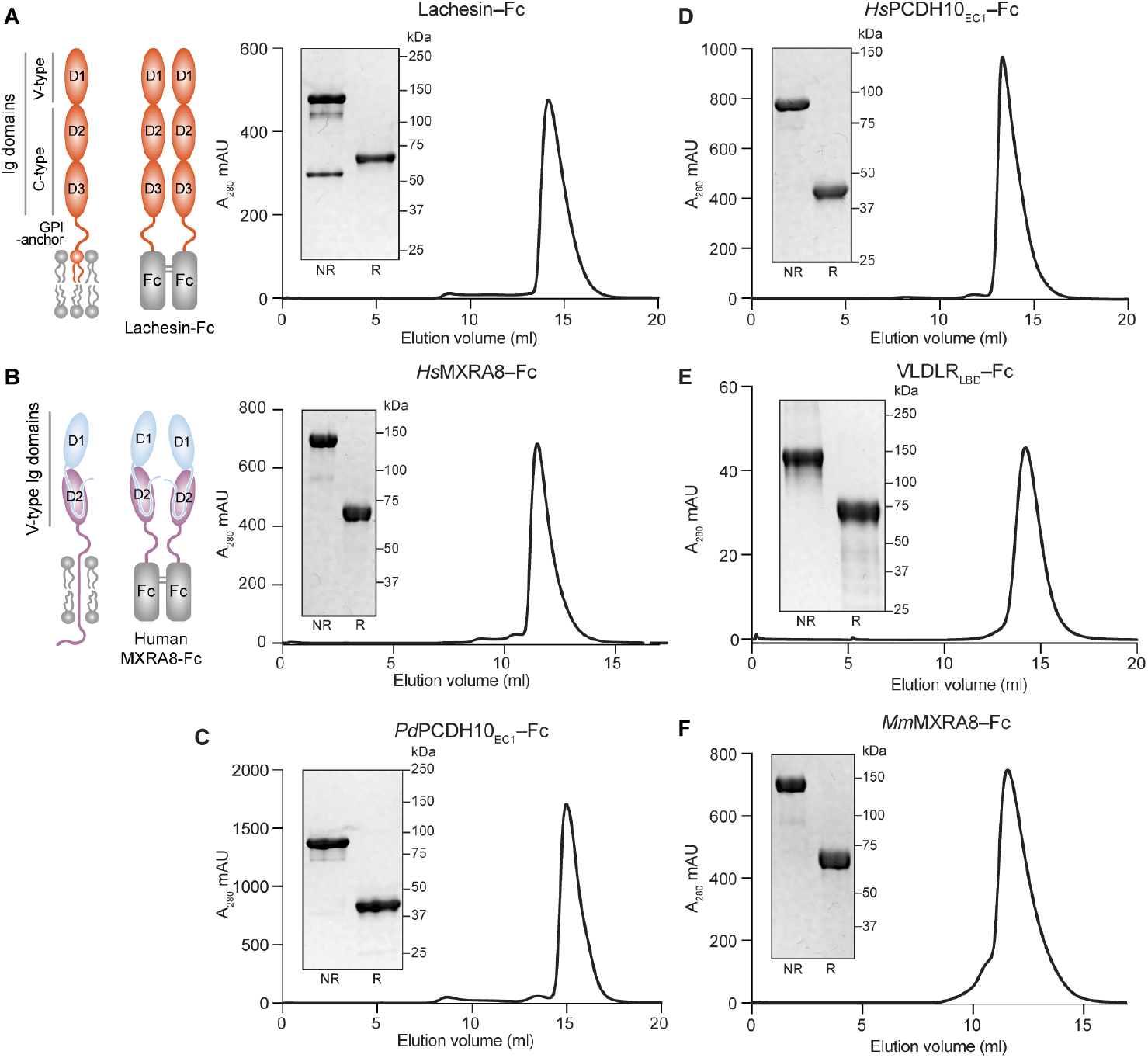
Purification of Fc fusion proteins. **(A)** Schematic diagram of Lachesin domain organization and Lachesin–Fc and size exclusion chromatography (SEC) trace of purified *Ae. albopictus* Lachesin–Fc. Immunoglobulin domains, including the variable (V) and constant (C) type domains, are indicated. Inset for the SEC trace is a Coomassie-stained SDS-PAGE gel of a peak fraction sample performed under non-reducing (NR) or reducing (R) conditions. **(B)** Schematic diagram of MXRA8 and domain organization and human MXRA8–Fc and SEC trace of purified human MXRA8–Fc. Inset is a Coomassie-stained SDS-PAGE gel. **(C–F)** SEC trace *Passer domesticus* PCDH10EC1–Fc (**C**), human PCDH10EC1–Fc (**D**), human VLDLRLBD–Fc (**E**), and *Mus musculus* (*Mm*)MXRA8–Fc (**F**). Inset is a Coomassie-stained SDS-PAGE gel.

**Figure S9.**
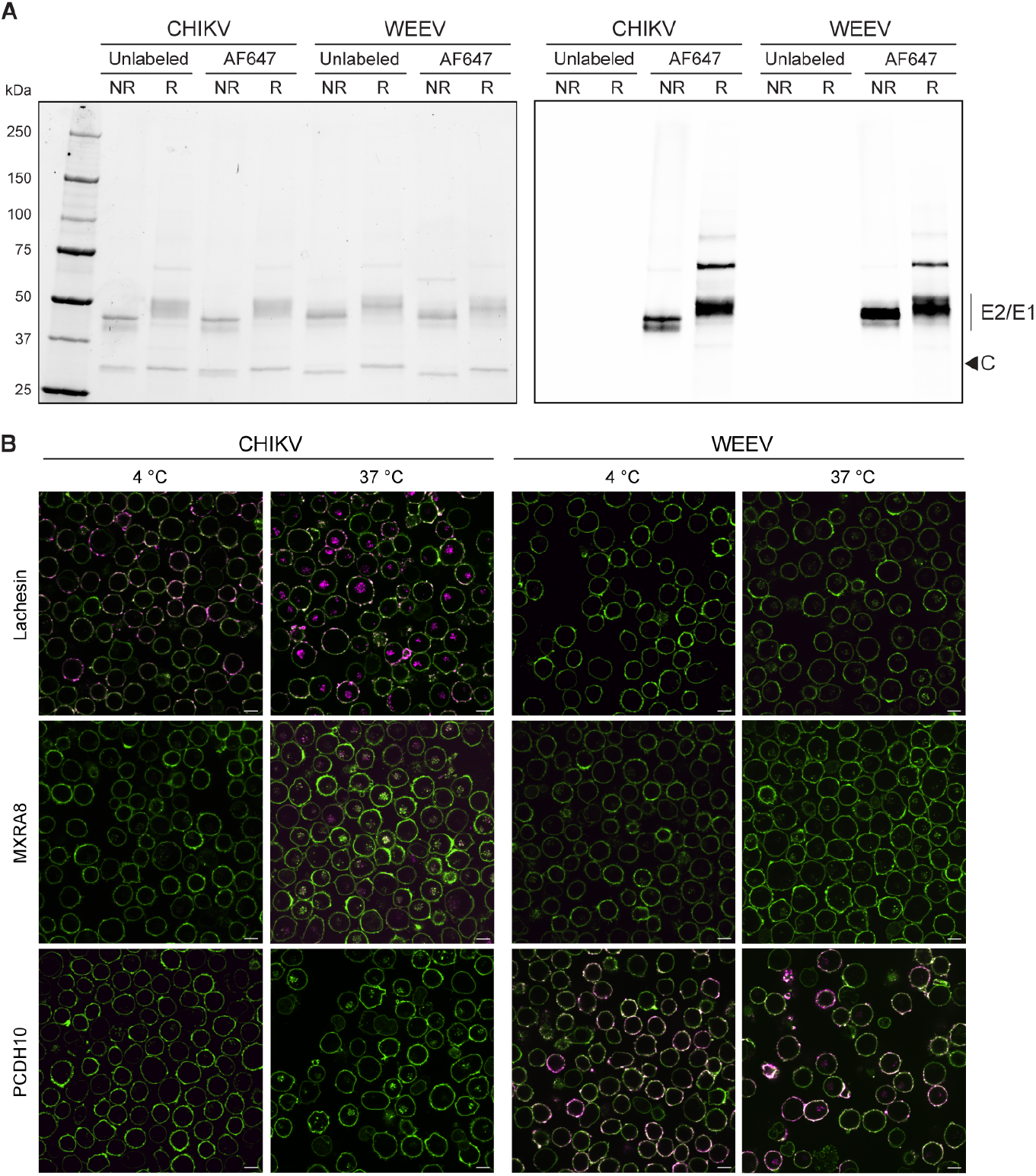
Gels of purified and labeled VLPs and confocal images of VLP attachment and internalization assays. **(A)** SDS-PAGE gel of unlabeled or AF647-labeled CHIKV and WEEV VLPs, visualized using a stain-free system (left) or lasers to excite the fluorophore (right). Capsid (C) and E2-E1 glycoproteins are denoted. Lack of fluorophore labeling of the capsid indicates intact VLPs. NR, nonreducing R, reducing. Representative gel of two separate experiments. **(B)** K562 cells stably expressing *Ae. albopictus* Lachesin, human MXRA8, or human PCDH10 were incubated with fluorescently labeled CHIKV or WEEV for 30 min at 4°C or 37 °C, stained with wheat germ agglutinin (WGA) and imaged by live cell confocal microscopy. Scale bar is 10 μm. Experiment was performed twice, and representative images are shown

**Figure S10.**
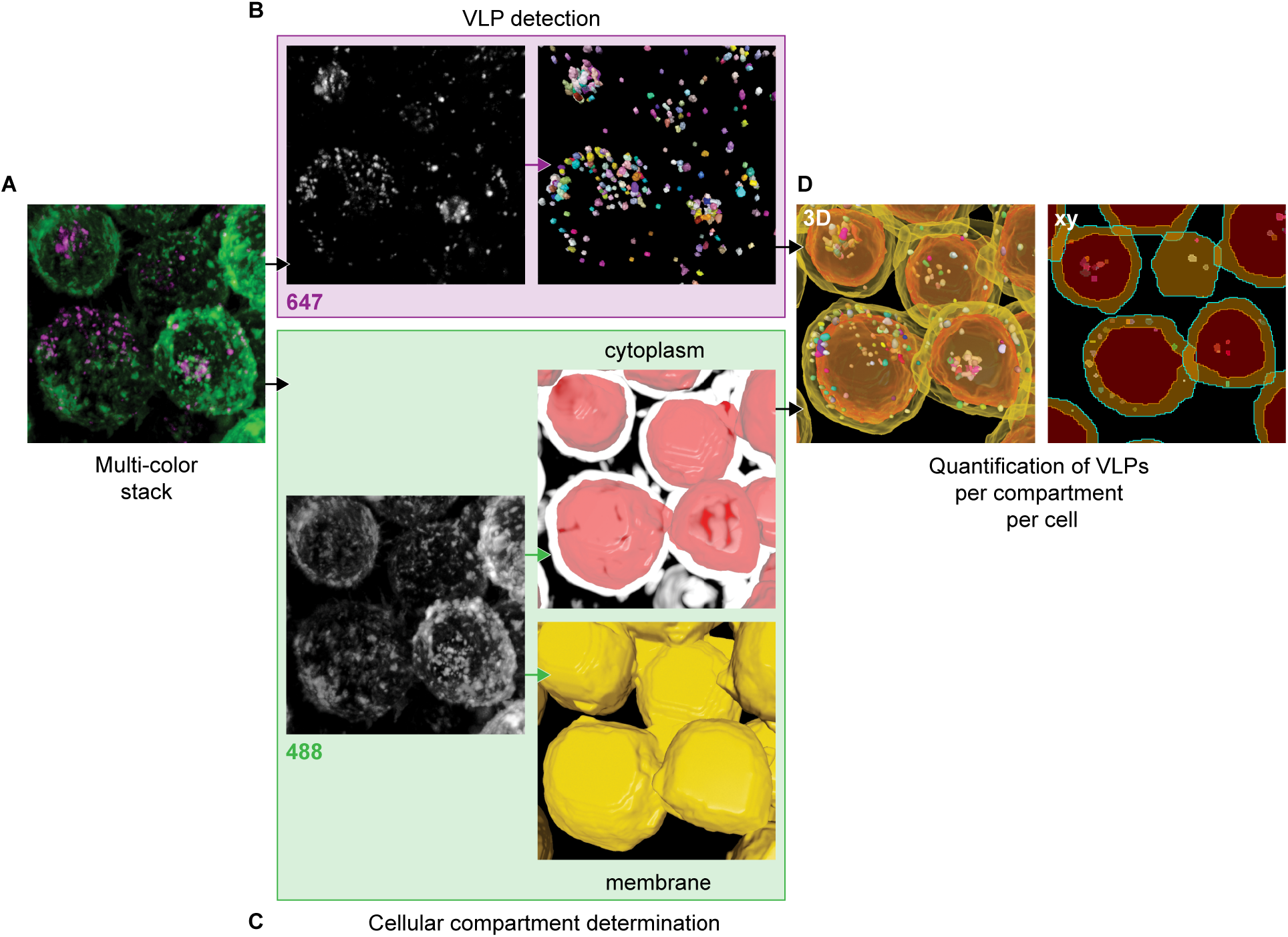
Workflow diagram of VLP attachment and internalization quantification. **(A)** 3D rendering of multi-color stacks (magenta, VLPs; green, cell membranes) using Arivis 4DFusion. Two custom-made pipelines were used to detect VLPs and cellular compartments. **(B)** 3D rendering of VLP stacks (left) and 3D rendering of detected VLPs (right). **(C)** 3D rendering of cellular membranes stacks (left), 3D rendering of the detected cytoplasm (red) overlayed with an enhanced-membrane filter (white) (right, top), and 3D rendering of the detected membranes (yellow) (right, bottom). The objects from each pipeline were pooled to count VLPs in each cellular compartment. **(D)** 3D rendering (left) or single slice of stack (right) of detected objects from the two pipelines combined, used to quantify the number of VLPs in each cellular compartment.

**Figure S11.**
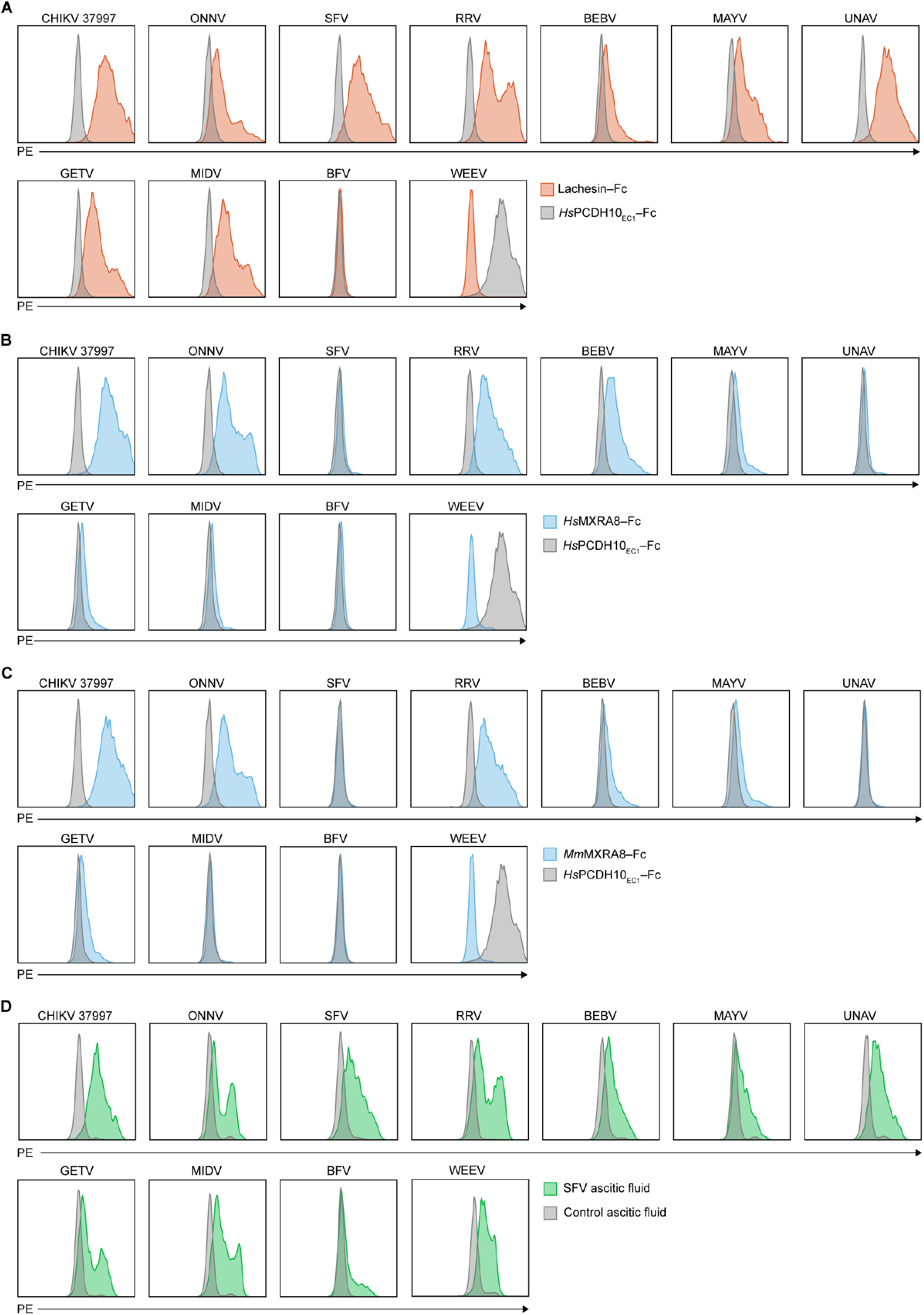
Fc fusion protein staining of cells expressing alphavirus E2–E1 glycoproteins. **(A–D)** Cell surface immunostaining of HEK 293T cells transfected with alphavirus E3–E2–(6K/TF)–E1 proteins of the indicated alphavirus were stained with 100 µg ml^-1^ *Ae. albopictus* Lachesin-Fc (**A**), human MXRA8–Fc (**B**), murine MXRA8–Fc (**C**), or mouse anti-SFV immune ascitic fluid (**D**). Glycoprotein sequences for the following strains were used: CHIKV 37997, ONNV SG650, SFV SFV4, RRV T48, BEBV MM2354, MAYV LET-1430, UNAV BeAr13136, GETV M1, MIDV MIDV857, BFV K61404, WEEV 71V1658. Representative plots shown for an experiment that was performed twice.

**Figure S12.**
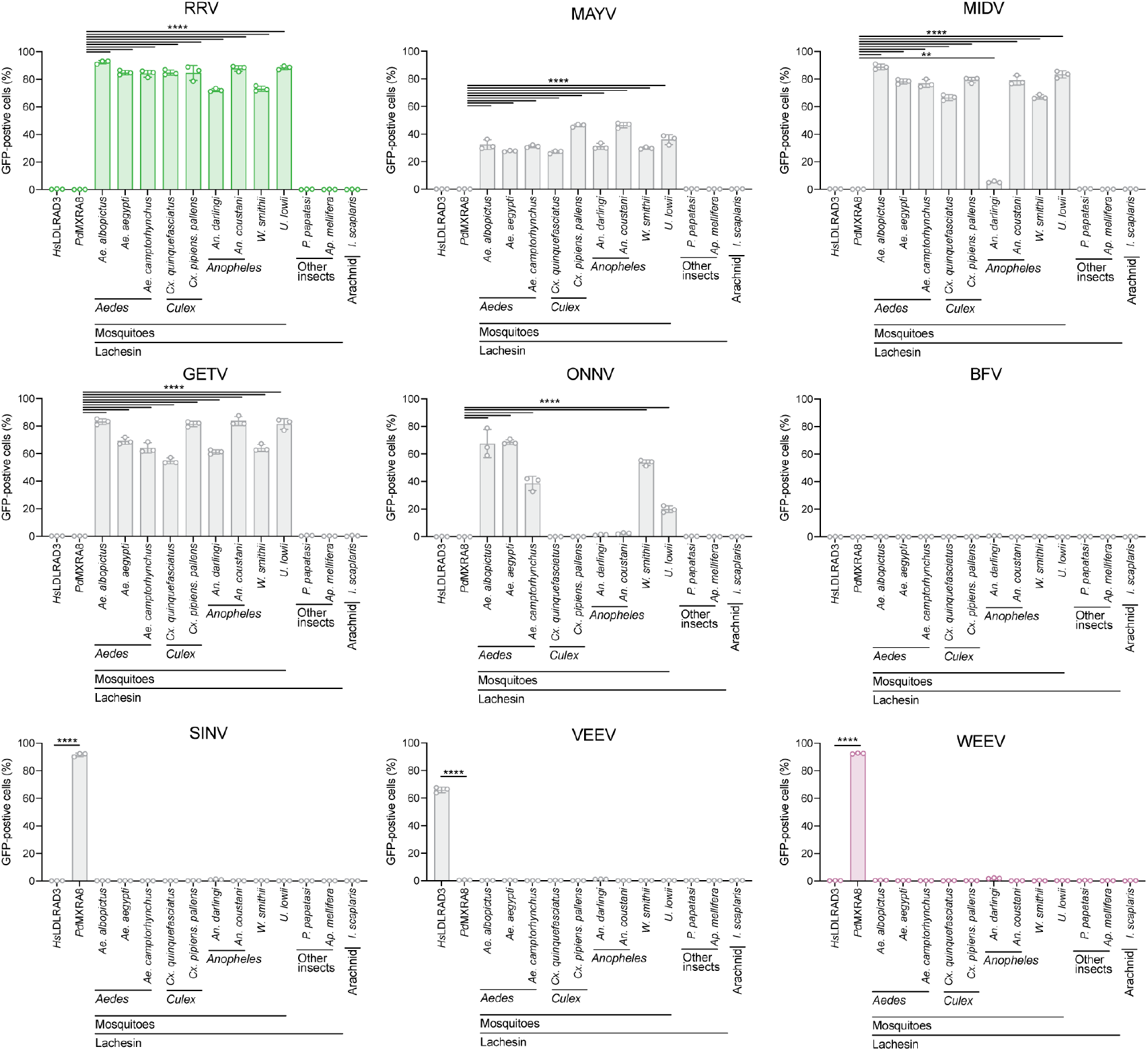
Alphavirus infections of cells expressing diverse Lachesin orthologs. K562 cells expressing Lachesin orthologs were infected with RVPs for RRV (T48), GETV (M1), MIDV (MIDV857) at an MOI of 2 as measure of K562-*Ae. albopictus* Lachesin, or ONNV (SG650) at an MOI of1 as measured on K562 cells expressing *Ae. albopictus* Lachesin, or MAYV (LET-1430) at an MOI of 0.5 measured on K562 cells expressing *Ae. albopictus* Lachesin, or WEEV (McMillan), SINV (TR339), VEEV (INH-9813) at an MOI of 2 measured on Vero E6, or BFV (K61404) at MOI = 3 measured on Vero E6. Infection as measured using flow cytometry 24 h post-infection. Data are mean ± s.d. of 3 experiments performed in duplicate (*n* = 3). Two-way ANOVA with Dunnett’s multiple comparison, \*\**P* = 0.0066, \*\*\*\**P* < 0.0001.

